# What We (Don’t) Know about Parrot Welfare: A Systematic Literature Review

**DOI:** 10.1101/2024.03.27.586789

**Authors:** Andrea Piseddu, Yvonne R. A. van Zeeland, Jean-Loup Rault

## Abstract

Parrots are popular companion animals but show prevalent and at times severe welfare issues. Nonetheless, there are no scientific tools available to assess parrot welfare. The aim of this systematic review was to identify valid and feasible outcome measures that could be used as welfare indicators for companion parrots. From 1848 peer-reviewed studies retrieved, 98 met our inclusion and exclusion criteria (e.g. experimental studies, captive parrots). For each outcome collected, validity was assessed based on the statistical significance reported by the authors, as other validity parameters were rarely available for evaluation. Feasibility was assigned by considering the need for specific instruments, veterinary-level expertise or handling the parrot. A total of 1512 outcomes were evaluated, of which 572 had a significant p-value and were considered feasible. These included changes in behaviour (e.g. activity level, social interactions, exploration), body measurements (e.g. body weight, plumage condition) and abnormal behaviours, amongst others. However, a high risk of bias undermined the internal validity of these outcomes. Moreover, a strong taxonomic bias, a predominance of studies on parrots in laboratories, and an underrepresentation of companion parrots jeopardized their external validity. These results provide a promising starting point for validating a set of welfare indicators in parrots.

## Introduction

Parrots have always fascinated human beings, influencing art, literature, religion across centuries and continents (1). Today, these birds, belonging to the order *Psittaciformes* (2), are one of the most popular companion animals after dogs and cats (3-6). Such popularity can be attributed to their bright and colourful plumages, but more importantly to their unique learning abilities which are comparable to those of human toddlers (7-10). Parrots, supported by large and neuron-rich forebrains (11, 12), can use and even manufacture tools (13-16), think economically (17, 18), succeed in problem solving, reasoning and planning tasks (19), and even remember their own past actions (20), an important prerequisite for self-awareness. Parrots also show different kinds of social competences: they can cooperate during problem-solving tasks (21-24), learn from conspecifics (25, 26) and exhibit prosocial behaviours (27-29). Similar to humans and a few other species, they can learn and imitate sounds (9, 30), and synchronize their motor output on incoming rhythmic acoustic or visual information (31-33).

These characteristics render parrots valuable and desirable companion animals, capable of positively fulfilling their caretakers’ cognitive and esteem needs (3, 34). However, such social and cognitive complexity make companion parrots especially prone to developing serious welfare issues when kept under inappropriate husbandry and management conditions. Examples of such welfare problems include higher risk of contracting viral, fungal or bacterial infections (e.g. avian bornavirus infection, aspergillosis, chlamydiosis), nutritional deficiencies (e.g. hypocalcaemia, hypovitaminosis A), and associated pathologies (e.g. metabolic bone disease, egg binding), as well as other non-infectious diseases (e.g. atherosclerosis, obesity), and development of fear-related, aggressive, stereotypic and/or self-injurious behaviours such as feather damaging behaviour (4, 35-37).

Experts from 51 different countries predict an increase in the trading of parrots due to their popularity (38). This will likely have detrimental consequences for the conservation of this highly threatened taxon (39, 40), but it also implies an increase of companion parrots population. Despite this prediction and the well-known welfare challenges of keeping captive parrots, there are currently no standardized guidelines for evaluating companion parrot welfare. The establishment of a parrot welfare assessment tool would represent a significant advancement in companion parrot welfare as it would facilitate welfare monitoring and could enhance the caregivers’ understanding of parrots’ needs, resulting in an improved quality of life. Suitable welfare indicators need to be identified to create such a tool and this can be achieved following five key steps.

The first step is to find and collect information from peer-reviewed scientific studies. Integrating scientific information represents the most appropriate strategy, as it allows welfare assessors to employ standardized and objective methods, lowering the risk of making assessments biased by personal experience, mood, and emotional subjective states (41, 42).

However, research findings may not always be valid and might represent inaccurate information (43, 44). Systematic reviews and simulations have shown that, due to weak experimental designs and settings, single research studies carry a high risk of bias, which is defined as “a systematic error or deviation from the truth, in results or inferences” (45). For this reason, the second step to identify welfare indicators is to verify the internal validity of the scientific findings collected. Internal validity is defined as “the extent to which the design and conduct of a study are likely to have prevented bias” (46), and it is typically divided in four sub-categories: construct validity (i.e. the extent to which a test measures what it is intended to measure) (47), face validity (i.e. appropriateness of the test and its parameters) (48), content validity (i.e. the extent to which the test covers the entire construct) (49), and criterion validity (i.e. the extent to which the outcomes of the test aligns with those previously obtained with validated instruments or “gold standards”) (50). “Internal validity” and “risk of bias” are closely associated (51); in fact, when a test presents high risk of bias, its results cannot be considered internally valid. Similarly, reliability, which is defined as production of consistent results within the same subject (*test-retest reliability*) (48), or between (*inter-observer reliability*) and within (*intra-observer reliability*) observers (52), is an important prerequisite for internal validity. A reliable measure is not necessarily valid; however, when the measure is not reliable, it cannot be valid (48). As such, it is important to screen scientific studies according to the aforementioned parameters to assess their internal validity.

The third step is to verify the studies’ external validity, i.e. the extent to which the findings of a study can be generalized and applied to other species, environmental conditions, or experimental settings (53, 54). This is especially important in the case of parrots as the *Psittaciformes* order comprises a vast diversity of species. Unlike “dog”, “cat” or “rabbit”, “parrot” is a general term grouping more than 350 species that are adapted to different ecological niches and have distinct environmental, dietary, and behavioural needs (55). This raises the question whether conclusions drawn from studies on a single species can be applied to other species (56). For our purposes, it is necessary to determine whether and to what extent the species of interest, i.e. those commonly kept as companion animals, have been studied. Similarly, the setting in which the study results have been obtained should be considered as living conditions in a zoo, shelter or laboratory differ markedly from those in a private household, thereby implying that results obtained under these circumstances are not necessarily applicable to parrots kept as companions.

The fourth step in the creation of an animal welfare assessment tool is to identify feasible measurements. As suggested by Yon and colleagues (57), animal welfare assessments should ideally be “rapid, non-invasive and should not require any specialist equipment, facilities or specific training”. This considerably reduces the risk of errors due to, for instance, assessors’ tiredness, instruments’ accuracy and sensitivity, or the animal’s reaction in response to handling.

According to Fraser (58), animal welfare should be assessed by employing measurements that reflect three distinct inextricable conceptual frameworks: the animal’s affective state, its biological functioning, and natural living. As such, the fifth and final step to create a welfare assessment tool requires capturing the various welfare dimensions through different indicators. Although behavioural indicators are considered the best reflection of animals’ ability to cope with their environment (56), a more accurate welfare assessment can be obtained by combining behaviour with other measurements, including physical condition, physiological parameters, presence of disease and pathologies, husbandry, nutrition, and management considerations.

Given the current lack of science-based welfare tools to evaluate the welfare of captive parrots, we conducted a systematic literature review in which we reframed the five key steps according to the following research questions: (i) Which, if any, scientific results related to the welfare of captive parrots can be considered valid and feasible welfare indicators? (ii) How many and which types of welfare indicators have been identified? (iii) From how many and which parrot taxa have these indicators been collected? and (iv) How much and what type of information is available specifically regarding companion parrots? Although our main target were companion parrots, we also collected information gathered from studies focused on other types of captive parrots, with the objective of identifying welfare indicators still applicable to our category of interest.

## Methods

All phases of this study were conducted following the PRISMA 2020 statement for reporting systematic reviews (59).

### Systematic Search

A systematic search was conducted to find all scientific studies relevant to the research questions. The population, intervention, control, outcomes (‘PICO’) strategy (60) was followed as much as possible and allowed to identify key terms related to the population of interest, to the type of intervention, and to the outcomes collected (note that the ‘control’ was not included as a search term as we did not restrict our search to case-control studies focussing on comparison of two interventions or comparison of the intervention with a control). We used terms such as “parrot”, “parakeet”, “psittacids” and the specific type of parrot (e.g. macaw, grey parrot) for the population; terms related to nutrition, husbandry and management (e.g. “foraging enrichment”, “diet”, “hand-rearing”) for the intervention; and terms such as “abnormal behaviour”, “disease”, “life span”, “emotional state”, or specific behaviour problems, diseases or pathologies (e.g. “feather picking”, “atherosclerosis”, “obesity”) for the outcome (see Table S1 for the complete list of the 86 search key terms). These key terms were then combined to create a search query using the Boolean operators AND, OR and NOT, and the queries were uploaded on the advanced search tool of the databases PubMed, CAB Direct and Web of Science (on May 16^th^, 2022).

### Paper selection: title and abstract screen, full-text screen

Following the literature search, the scientific studies found were uploaded into a reference manager (Endnote X7; (61)). After removal of duplicates and triplicates, all remaining texts were screened twice using exclusion and inclusion criteria (Table S2). Studies could involve parrots from any species, gender or age, and a variety of different interventions and outcomes (as specified in Table S2) and were included for further evaluation as long as the study was conducted in a captive population and was not focused on reproductive parameters (egg hatchability, number of eggs) or chick development as we considered these irrelevant for companion parrots. Further restrictions were related to language (English only), methodology (no studies involving <5 subjects or without statistical analysis), publication type (full papers describing original research only) and retrievability of the paper. The first screening, based on the title and abstract, was conducted by one reviewer (AP), and aimed to exclude all studies that were considered irrelevant to address the research questions. All studies that passed this initial screening subsequently underwent a second screening, in which the full-text was read by two independent reviewers (AP, J-LR, see supplemental materials). Additional studies found through external sources (e.g. cited references) were also considered for their eligibility to ensure to encompass the latest literature available (updated till January 27^th^, 2023).

### Data collection

#### Collection of the outcome measures and corresponding risk factors

All behavioural, physiological, physical and health parameters that could potentially be linked to parrot welfare were collected from studies that passed the exclusion and inclusion criteria. Only parameters that were included as ‘outcome measures’ (62), i.e. as part of the experimental design, were considered for further evaluation. For instance, outcome measures such as feeding behaviour or body weight were collected and considered only when measured specifically in relation to examined risk factors (e.g. social isolation, unbalanced diet, cage size or use of enrichments) but not when correlated with natural biological phenomena (e.g. breeding season changes). If not explicitly written by the authors, the potential risk factors were interpreted from the experimental conditions. For instance, in studies that compared behaviours of enriched and non-enriched parrots, the main risk factor for welfare was the lack of enrichment, whereas in studies comparing the behaviours of hand-reared versus parent-reared subjects, the risk factor was the rearing method. All risk factors associated with outcome measures with a p-value <0.05 (both feasible and non-feasible) that were easy to define and identify (e.g. social isolation, diet, cage size etc.) were collected.

#### Internal validity and feasibility of outcome measures

Multiple outcome measures could be identified from a single study, and because our focus was on identifying outcome measures that could be used as parrot welfare indicators, we assessed the internal validity of each outcome measure separately, to the difference of other systematic reviews that did it for the study as a whole. The risk of bias was assessed by employing selected domains of the SYRCLE’s RoB tool (63): from the main text, we extrapolated whether outcomes measures were statistically linked to risk factors (p-value significance), whether data were measured and collected by a blinded assessor, whether studied population was randomized (where applicable), and, in case of experimental setups with multiple conditions, whether the design was balanced between subjects and/or groups. In addition, we checked for tests of intra- and inter-observer reliability (see Table S3 for definitions of each of the evaluated validity parameters).

Given that we aimed to find welfare indicators that could be used to assess parrot welfare in practice, we also evaluated the outcome measures for their feasibility. They were classified as feasible if the behavioural assessment or measurements could be readily performed, requiring only the use of commonly available equipment (e.g. weight scale) or the use of minimally-invasive routine handling techniques, or as non-feasible if the use of specific instrumentation, calculation or veterinarian-level skills or expertise were required.

#### Grouping of outcome measures in welfare categories and welfare dimensions

To facilitate interpretability of the results and determine how many outcome measures of a certain type could be identified, the outcomes were grouped in categories according to commonalities in their underlying biological construct (e.g. stereotypic behaviour, body condition, foraging behaviour) and subsequently classified according to one of eight distinct welfare dimensions (i.e. physical or physiological measure, abnormal and fear-related behaviours, maintenance behaviours, locomotory behaviours, exploratory and foraging behaviours, social behaviours, and diseases and pathologic conditions; Table S4).

#### Extrapolability of outcome measures across species and settings

To determine the extent to which outcome measures would be applicable or extrapolable to other species or settings, we collected taxonomic data (parrot species and genus), where available. Data acquired from multiple genera was classified under the category “multiple”. Following the definition of “pet” or “companion” animal proposed by Farnworth (64), the studied subjects were identified as companion parrots when they “lived with humans or within human social structures where they were provided with some, or all, of their needs” and “played a primarily social role within a household or community”. Alternatively, we identified the parrots based on their living conditions as parrots kept in laboratories, zoos, shelters or breeding centres (see Table S5 for the complete list and definitions of subjects’ living conditions).

#### Data analysis

The initial aim was to assess the risk of bias of all possible identified welfare indicators using the validation parameters described above (Table S3). However, as most papers lacked information regarding intra- and inter-reliability and blinding of the study design, we consequently decided to only partially assess their internal validity considering only statistical significance (alpha threshold fixed at 0.05), excluding all outcomes with a p-value ≥ 0.05, as measures with higher p-values were more likely to represent information that could not reliably be linked to a given risk factor. Outcomes with a significant p-value were subsequently screened for their feasibility. The outcome measures with a significant p-value and considered feasible were then grouped according to the welfare categories and dimensions they belonged, and according to subjects’ characteristics (genus and living condition). Data was analysed using the R statistical software (65) and the R package “dplyr” (66). Figures were created using the R package “ggplot2” (67).

## Results

### Result of the systematic search and paper selection

The systematic search led to the collection of 1946 scientific studies: 697 from CAB direct, 657 from Web of Science and 592 from PubMed. A total of 189 studies were found to be duplicates and triplicates, and, after removing these, the total amount of hits dropped to 1848. The first screening, based on title and abstract reading, led to the selection of 140 studies. The second screening, based on full-text reading, led to the retention of 83 studies. An additional 15 studies were found from external sources. This screening step led to a final amount of 98 studies from which data was collected (Figure 1).

**Figure 1.**
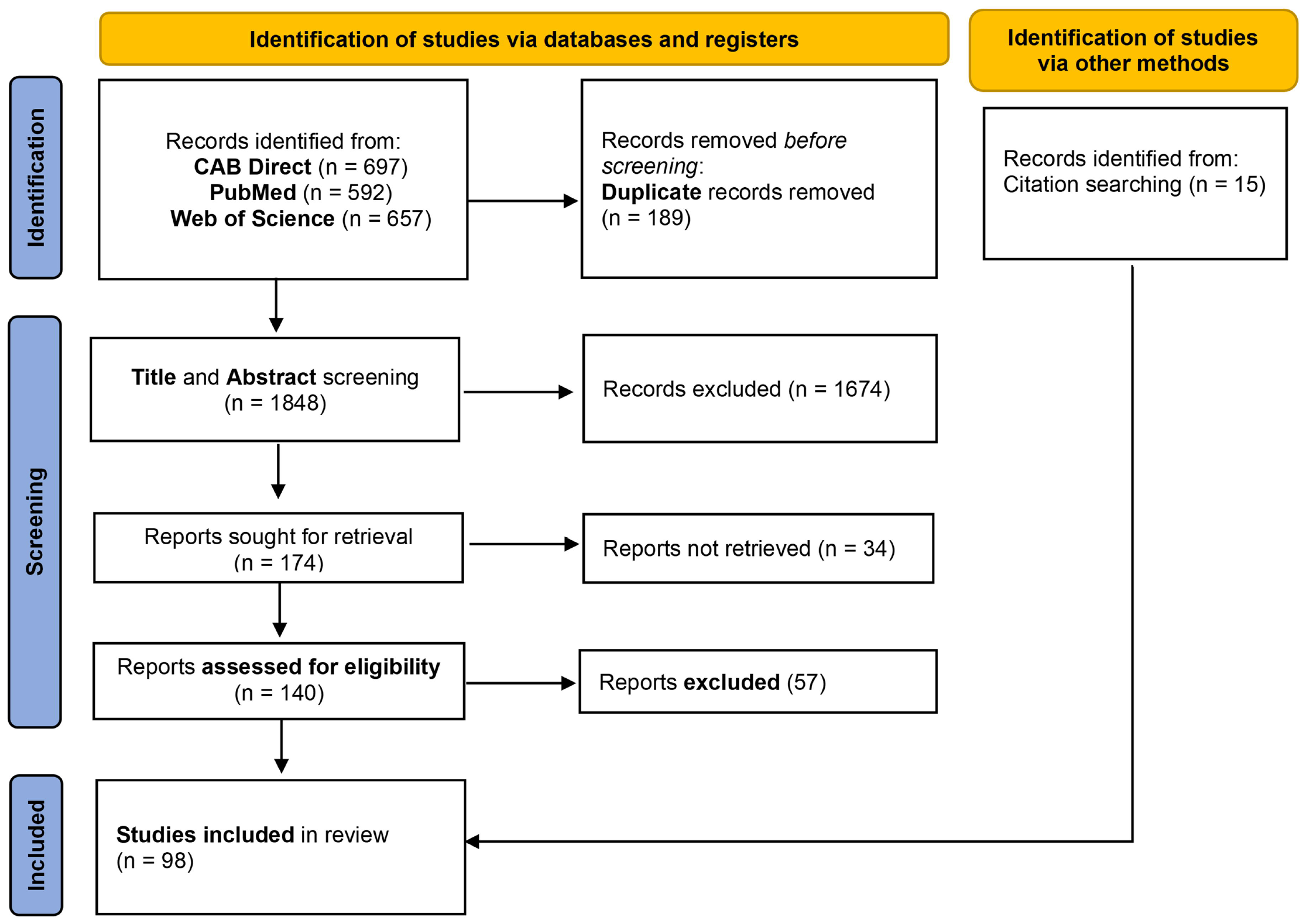
PRISMA Flowchart. The PRISMA flow diagram for the systematic review showing the studies identified from each database, number of studies screened, excluded, and included.

### General results

The year of publication of eligible studies ranged from 1992 to 2023, with 76 studies (77.6%) published between 2010 and 2023. Of the studies that were eligible for full evaluation, 95 (96.9%) reported significant results and, of these, 72 (73.5%) reported outcome measures that were considered feasible to be carried out by owners (see Table S8). Only 13 of these 72 studies with significant and feasible outcomes (13.3%) specifically related to companion parrots (see Table S8). The number of outcomes collected in a study ranged from 1 to 100 (median: 11). Of the total of 1512 outcomes collected, 720 (47.6%) had a significant p-value, and of those 572 (37.8%) were also considered feasible. Of these 572 outcomes, 68 (4.49%) were obtained from companion parrots.

### Risk of bias

Intra and inter-observer reliability and assessor blindness were reported for less than 5% of the outcome measures and were not specified for more than 77% of the outcomes (Table 1). In addition, a high number of outcome measures were from studies that, due to their experimental set up, did not allow to control for the presence of biases or prevent it. For instance, outcomes collected from questionnaires and retrospective studies (17.5 %) could not be tested for intra- and inter-observer reliability, and random assignment of subjects to groups, assessor blindness and balancing of experimental conditions could not be applied to this type of studies. Another example came from studies where subjects were assigned to control and enriched groups: in this case, it may not have been possible to blind the assessors and therefore to control for this specific bias. Due to these circumstances, we could not establish with certainty which outcomes measures could be classified as valid welfare indicators.

**Table 1.** Assessment of the risk of bias. The table shows the levels of reporting for each validity parameter, as the proportion of all outcome measures (%).

| Reporting | Validity parameters |  |  |  |  |
| --- | --- | --- | --- | --- | --- |
|  | Inter-observer reliability | Intra-observer reliability | Random assignment of subjects to groups | Assessor blinded | Balancing of experimental conditions |
| Yes | 2.9% | 3.4% | 17.9% | 2.2% | 15.7% |
| No | 0% | 0% | 0% | 1.1% | 0.1% |
| Not specified | 77.5% | 77.1% | 3.8% | 33.1% | 16.9% |
| Not possible | 19.6% | 19.5% | 78.3% | 63.6% | 67.3% |

### Representation of welfare dimensions and categories

#### Outcomes classified according to welfare dimensions

Out of the 8 welfare dimensions, the welfare dimension with the highest number of feasible and significant outcomes was “social behaviours” (n=141), whereas for most other dimensions the number of significant and feasible outcomes ranged between 80 and 93 (Figure 2 and Table S9). The welfare dimension “physiological parameters” included a high number of significant outcomes (n=97); however, all of these were considered not feasible as they required invasive sampling techniques (e.g. venipuncture) and laboratory equipment. A similar trend was noted for the welfare dimension “diseases and pathologic conditions”, albeit the number of outcomes reported was lower to start with (Figure 2).

**Figure 2.**
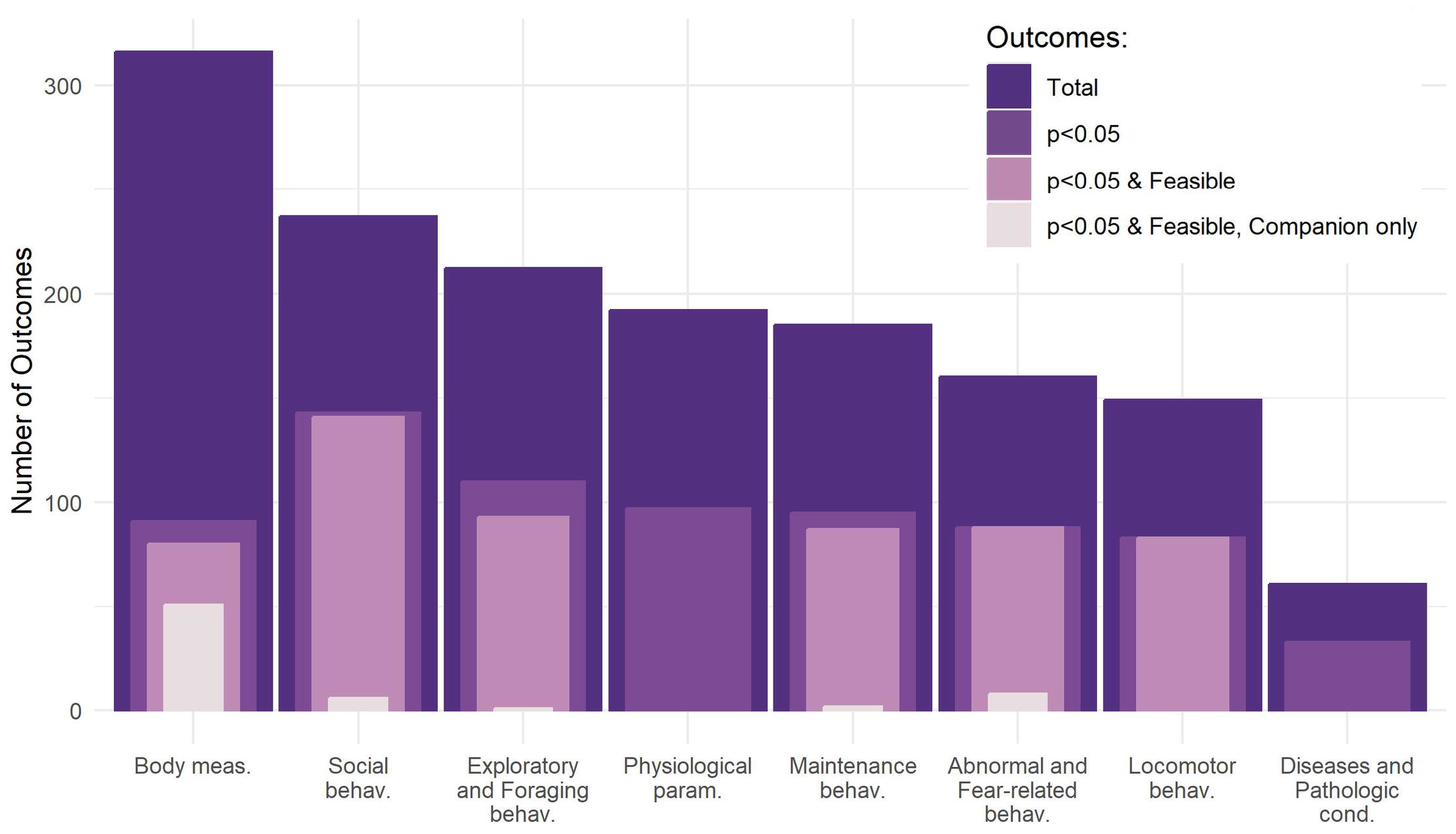
Number of outcomes grouped by welfare dimensions. The overlapped bar plot shows on the y-axis the number of total outcomes collected, significant outcomes, significant and feasible outcomes, significant and feasible outcomes collected from companion parrots, grouped by welfare dimensions. Behav.=behaviour, meas.=measurements, param.=parameters, cond.=conditions.

#### Outcome measures classified according to categories

The significant and feasible outcome measures were grouped in 26 different welfare categories (Table S10). “Stereotypies” covered the highest number of significant and feasible outcomes measures (n=77) and captured oral stereotypies (e.g. wire or sham chewing, beak grinding), head stereotypies (e.g. spot pecking), and locomotor stereotypies (e.g. pacing, route tracing). “Indirect measures of feather-damaging behaviour” included multiple scoring methods related to feather condition or feather improvement and was the category with the second highest number of significant and feasible outcome measures (n=69), followed by “self-care” (n=42) with behaviours such as preening and stretching (Table S10). “Human-animal interaction” was the category with the highest variety and included measures such as response to unfamiliar and familiar handlers, human-direct aggressiveness, and food acceptance (Table S10). Outcome measures such as body weight and body mass were grouped in category “body condition”; walking, climbing, and flying in “locomotion”; food intake and feeding bout in “feeding”; crown, nape, cheek feather ruffling and crest erection in “facial and body displays”. Of the 68 significant and feasible outcomes measures collected from companion parrots, 51 referred to feather damaging behaviour, 5 to human-animal interactions, 5 to stereotypies, 3 to fear-related behaviour, 1 to foraging behaviour, and 1 to sexual behaviour (Table S12).

### Association between risk factors and outcomes measures

Significant and not feasible outcome measures were grouped in 8 welfare categories and were related to various risk factors (Table S11). For instance, human (neonatal) handling was linked to changes in the immune system (68), increased respiration rate (69) and in serum corticosterone concentrations (68); social isolation affected telomere length (70); indoor housing and lack of UV-B lighting increased the risk for vitamin D deficiency (71, 72). An unbalanced diet was correlated with changes in several parameters: feather colour (73), immune system responses (73, 74), plasma, aortic, arterial and hepatic cholesterol levels (75) as well as echocardiographic parameters that were considered to be associated with cardiovascular dysfunctions (76), and higher incidence of atherosclerosis (74) (Table S11).

Risk factors were also identified for significant and feasible outcome measures. For instance, the lack of environmental enrichment was associated with 18 welfare categories (Table S10). Social isolation represented a risk factor for developing stereotypies (77), and was associated with reduced preening (77), flying (78) and locomotor activities (79), and also with an increase in vocalizations (78) and avoidance behaviour towards humans (79). Rearing method was correlated with the development of feather damaging behaviour (80, 81), stereotypies (77) and preferential interactions with humans (80) (Table S10). Cage size was associated with the emergence of phobic behaviours (80), abnormal behaviours such as incessant screaming, oral and locomotor stereotypies, increase of courtship behaviours and singing (82), and changes in locomotor activities (walking, climbing) and preening (82, 83) (Table S10). Regarding the significant and feasible outcome measures collected from companion parrots, risk factors related to human-animal interactions were the most common, followed by housing characteristics (Table S12).

### Representation of the various genera and living condition

#### Living conditions represented in the studies

The majority of studies (n=48) were conducted on parrots kept in laboratories, followed by 22 on companion parrots, 10 on parrots kept in zoos, 6 on parrots kept in breeding facilities, 3 on parrots kept in rehabilitation centres and 2 on parrots kept in shelters (Table S6). Two studies focused on comparing parrots kept in different settings (i.e. wild versus companion parrots versus parrots kept in a zoos or breeding facility, and companion versus parrots kept in laboratories; Table S6). For 7 studies, it was not possible to define the living condition of the parrots (Table S6).

#### Outcome measures in relation to genera

Outcome measures were collected from 13 genera, of which 10 belonged to the superfamily *Psittaccoidea* and 3 to the superfamily *Cacatuoidea. Melopsittacus* and *Amazona* were the genera with the highest number of significant and feasible outcomes (respectively n=150 and n=128), followed by the *Ara* (n=72), *Nymphicus* (n=65), and *Psittacus* (n=54) genera (Figure 3 and table S8). Fifty-four (9.44%) significant and feasible outcome measures were available from studies that included multiple genera (see Table S7). For other genera, the number of significant and feasible outcomes collected ranged between 1 (*Guaruba*) and 17 (*Pyrrhura*). For the genus *Platycercus* no significant and feasible outcome measures were identified (Figure 3, Table S8). A total of 68 feasible and significant outcome measures were collected from companion parrots, with 32 (47%) outcomes related to multiple genera, 24 (35.4%) to the genus *Psittacus*, 9 (13.2%) to the genus *Cacatua*, and 3 (4.4%) to the genus *Agapornis* (Figure 3, Table S12).

**Figure 3.**
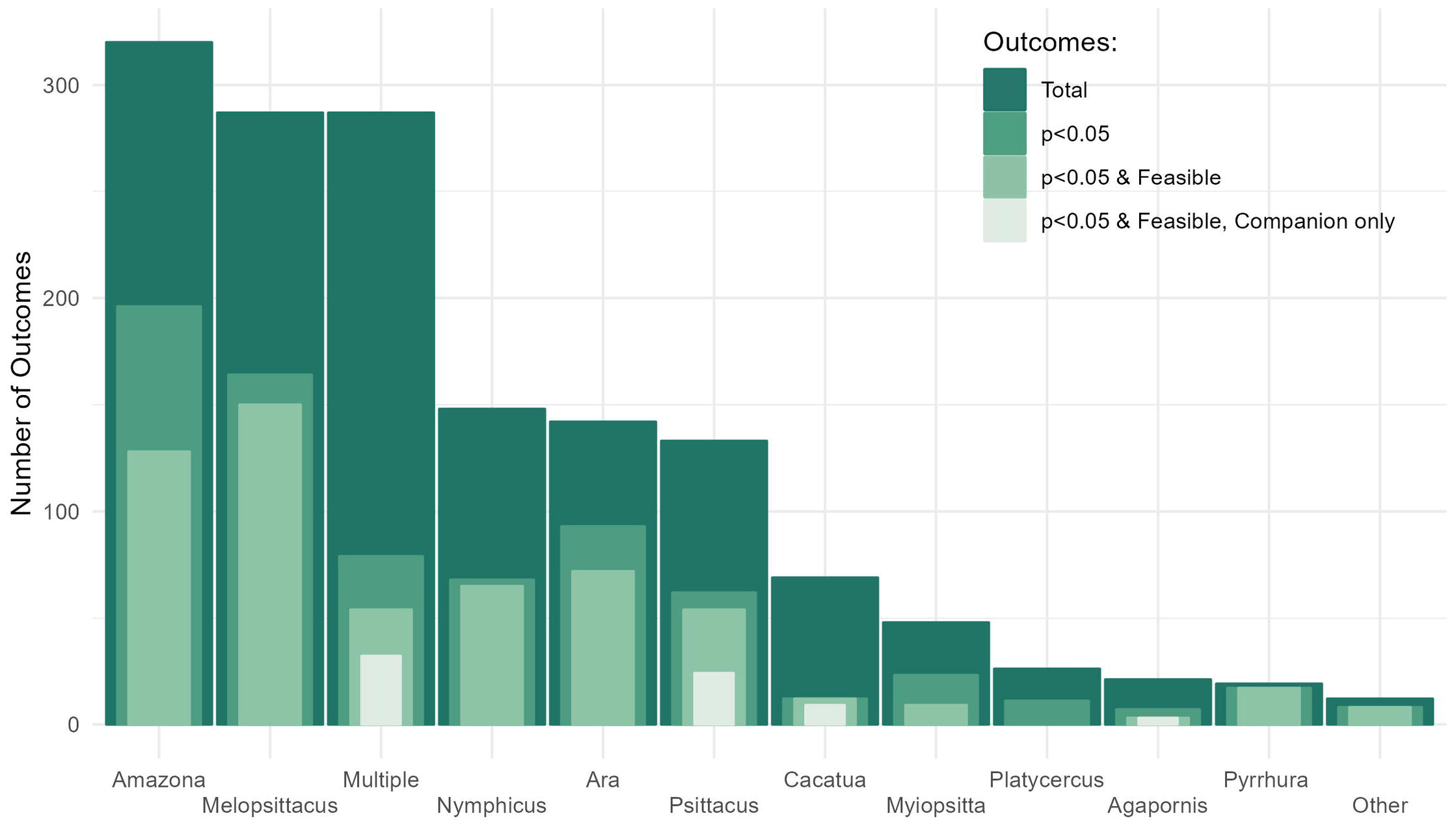
Number of outcomes grouped by parrot genera. The overlapped bar plot shows on the y-axis the number of total outcomes collected, significant outcomes, significant and feasible outcomes, significant and feasible outcomes obtained from companion parrots, grouped by parrot genera. “Other” refers to the pooled genera *Calyptorhynchus, Guaruba* and *Loriculus*.

#### Relationship between genera and welfare dimensions

Of the welfare dimensions for which we identified significant and feasible outcomes, “social behaviours” was the one investigated in the highest number of genera (9 out of 13). All other welfare dimensions covered 8 genera, except the dimensions “abnormal and fear-related behaviour” and “locomotor behaviour” which both were covered by 6 genera (Figure 4). The genera-welfare dimension association with the highest number of feasible and significant outcomes was the combination *Melopsittacus*-”abnormal and fear-related behaviours” (n=62), followed by *Nymphicus*-”social behaviours” (n=41), *Amazona*-”exploratory and foraging behaviours” (n=38), *Melopsittacus*-“maintenance behaviours” (n=35), and “Multiple genera”-”body measurements” (n=29) (Figure 4). Three genera were covered by only one welfare dimension: *Agapornis* and *Guaruba* with “body measurements”, and *Loriculus* with “exploratory and foraging behaviours” (Figure 4).

**Figure 4.**
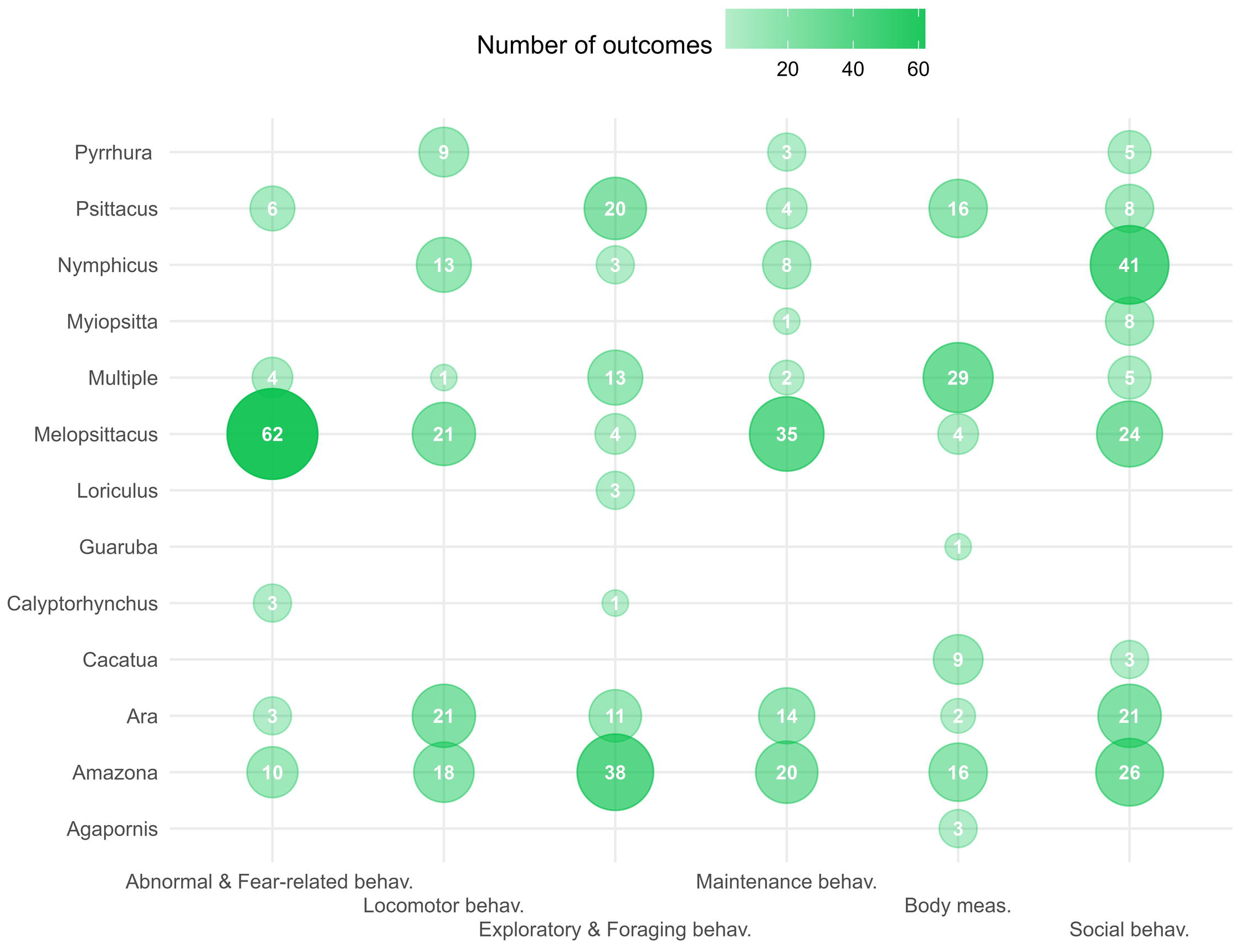
Bubble plot showing the number of significant and feasible outcome measures covered by each genus-welfare dimension combination. Welfare dimensions are reported on the x-axis and parrot genera on the y-axis. The number inside each bubble represents the number of significant and feasible outcomes belonging to a specific welfare dimension and collected from a specific genus. Increasing bubble size and opacity indicates a higher number of outcomes covered by the genus-welfare dimension combination. Behav.= behaviour, meas.=measurements.

## Discussion

### Internal validity

The main aim of this systematic study was to identify potential welfare indicators for companion parrots by first assessing the internal validity of outcome measures from published scientific studies. We found a high risk of bias associated with the outcome measures in the scientific literature. For instance, intra- and inter-observer reliabilities and assessor blindness were almost never reported. We thus only consider outcomes that presented a significant p-value. We identified 572 outcomes measures that presented a significant p-value and that we classified as feasible, but these need to be thoroughly validated before being used as welfare indicators in practical assessments.

### Welfare dimensions

The significant and feasible outcome measures linked to parrot welfare were well distributed across six of the eight welfare dimensions, except for physiological parameters, and disease and pathological conditions. Behaviours represented the most common type of outcomes measures, covering five out of eight welfare dimensions, and 24 out of 34 categories.

“Social behaviour” was the welfare dimension with the greatest variety of significant and feasible outcome measures and welfare categories. Within this dimension, we found several behaviours such as vocalization, mate-related behaviours, aggressiveness, allopreening, or behaviours related to human-animal interactions. The latter presented a remarkable heterogeneity of outcome measures, including several outcomes linked to inappropriate handling and physical contact. Human-animal relationship plays a fundamental role in guaranteeing companion animals’ positive welfare (84); for this reason, these results represent a good starting point for the development of an assessment tool tailored to companion parrots. Outcome measures of the welfare category “facial and body display” were the only results retrieved that proposed the use of facial expressions (feathers ruffling, blushing) as indicators of calmness and positive human-parrot interaction (85-87) and the display of erected crests as a sign of high arousal (88). Many others parrots’ displays such as body postures have been interpreted as ways parrots communicate their level of arousal or signal an imminent aggressive responses (89), however this information is not supported by experimental studies. Observing facial and body displays can be useful to assess welfare, for example in preventing negative interactions with caretakers or other animals that live in the same environment; however, further investigations are needed to validate such indicators.

Three welfare dimensions concentrated almost half of the significant and feasible outcomes: “locomotor behaviours”, in which we grouped behaviours such as flying and climbing; “exploratory and foraging behaviours”, reflecting the way in which parrots explore new environments and novel objects and interact with enrichments; and “maintenance behaviours”, in which we clustered behaviours such as feeding and resting. All these behaviours are used while foraging, an important activity for wild parrots as it occupies between 40% and 70% of their daily active time (90, 91). Therefore, indicators linked to foraging are suggested to be relevant to monitor captive parrot welfare.

For the welfare dimension “abnormal and fear-related behaviours” we retrieved several outcomes that could be used to assess welfare: incessant screaming, phobic behaviours, and many types of stereotypies such as locomotor (e.g. route trace, pacing), oral (e.g. wire chewing) and whole body (e.g. rocking, bobbing) stereotypies. Feather damaging behaviour, a common abnormal behaviour in companion parrots, especially in the genera *Psittacus* and *Cacatua*, has been the subject of many studies as it leads to several medical problems (92). However, feather damaging behaviour is difficult to observe directly, which may explain why we retrieved only one study where authors recorded duration and frequency of this self-injuring behaviour (93). Stereotypies and feather damaging behaviour, however, do not always reflect the current welfare state of the subject as they may also manifest themselves as “behavioural scars” and can remain even after the stressful stimulus or situation that triggered them is no longer present (94). For this reason, they should be included in a parrot welfare assessment scheme but accompanied by the corresponding risk factors identified, such as social isolation, lack of enrichment, or living in a small cage.

“Body measurements” was the only welfare dimension to include significant and feasible outcome measures that were not behaviours and consisted of two categories. One was “indirect measures of feather damaging behaviours” and included the outcomes “presence of feather damage (yes/no)” and plumage scores. These two outcomes can be used to detect the presence of feather damaging behaviour. Additionally, plumage scores allow caretakers to monitor improvement or deterioration of plumage condition over time. Moreover, previous studies showed good to excellent agreements within and between observers for this type of measurement (95, 96). It is important to highlight that damage to the plumage is not specific to behavioural disorders, as it can also be caused by other factors that are still relevant to parrot welfare such as malnutrition, virus infections, parasites infestation or inappropriate husbandry or management (e.g. small or overcrowded cage) (97). Due to their high feasibility and their well-established link to the welfare of captive parrots, “indirect measures of feather damaging behaviours” can be considered the most promising welfare indicators among all types of outcomes collected. The second welfare category in the dimension “body measurements” was “body condition, which contained measures indirectly linked to body fat composition, such as body weight, chest girth, and body mass. Several studies included in this systematic review tested the effect of an unbalanced diet on parrots’ body condition; however, most of the results were not significant. Caloric deficit and surplus are responsible for changes in body fat composition in most animal species, and, arguably, this also applies to parrots. As such, the lack of significant findings might be due to experimental design or the lack of sensitivity of the outcome measures collected. However, some studies found body fat composition to be influenced by regular physical activity (98, 99) and prolonged exposure to artificial light at night (100), two aspects that are seemingly understudied but could be highly relevant for companion parrot welfare.

We could not retrieve any feasible outcomes for the welfare dimension “physiological parameters”, mostly due to these requiring specialized techniques or equipment for collection or analysis. However, we identified several risk factors that can be associated with changes in these physiological parameters. Husbandry and management conditions were recurrent risk factors that influenced parrots’ physiology, especially stress-related parameters. For the welfare dimension “diseases and pathologic conditions”, which also lacked feasible outcome measures, risk factors mostly included demographic characteristics like parrot age, sex, or species. None of the studies retrieved looked for physical measurements as a clinical sign for existing disease or pathology. This finding was unexpected considering that these types of measurements are heavily influenced by health problems; for instance, upper beak and nail overgrowth, increased weight, and changes in feather colour and quality are anecdotally reported as signs indicative of (fatty) liver disease (101). Some of these parameters, commonly used by veterinarians based on expert knowledge or experience, were not reflected in the scientific literature or may have been missed in our search as it is difficult to comprehensively capture all possible health problems. Further experimental validation of some of these commonly used clinical diagnostic signs would be valuable.

### Most relevant risk factors for companion parrots

Among all risk factors associated with poor welfare, four emerged as especially important for the welfare of companion parrots. As suggested by several authors, captive parrots need to be mentally stimulated with different types of enrichment in order to prevent boredom, frustration, and other conditions associated with poor welfare (36, 102). In support of this, we found that a lack of physical and foraging enrichment was the most recurrent risk factor and was associated with changes in maintenance, locomotor, exploratory and social behaviours, and with expression of abnormal behaviours. A recent study on grey parrots (*Psittacus erithacus*), not included in our results, demonstrated that combining two different enrichment devices stimulated both the appetitive and consummatory phases of foraging behaviour, resulting in increased daily foraging time (103); hence not just the provision but also the design of enrichment devices is important. Moreover, other forms of enrichment that have been less investigated (e.g. cognitive and auditory) warrant research.

Social deprivation and social isolation also appeared to be recurrent risk factors in our results and were associated with outcome measures belonging to 7 out of the 8 welfare dimensions. Parrots are highly social species (104) but are often housed alone as companion animal, a living condition that we found being linked to poor welfare. A recent study showed that parrots living without other parrots were more likely to show problematic behaviours such as biting humans and stealing human food, and parrots left alone for more than 6 hours daily tended to be more prone to show feather damaging behaviour (105).

Personality also influences how parrots interact with their environment and cope with challenging situations. Several studies included in this systematic review showed that specific personality traits or coping styles were linked to the emergence of feather damaging behaviour (106), to the exhibition of attention bias (107), to the time spent feeding and interacting with the enrichment (108), and to fearful responses towards humans (109). Assessing personality can be an effective strategy for improving the welfare of captive animals (110), but to date there are few studies investigating how personality could impair parrot welfare.

Rearing methods also emerged as a crucial risk (developmental) factor that may impact parrots’ quality of life and welfare. Neonatal handling of parent-reared chicks resulted in reduced aggressiveness and fear-related and feather damaging behaviours in later life (68, 111), whereas hand-rearing was linked to these problematic behaviours (80, 81, 112). However, given that hand-rearing might induce irreversible changes, the results observed from the studies should be used with informative and preventive purpose, as these cannot be changed after weaning.

Overall, our data points to a lack of enrichment, social isolation, personality, and rearing method as important aspects for companion parrots that should be taken into account in parrot welfare assessment.

#### External validity

The second aim of this systematic review was to assess the external validity of the outcomes by establishing from which species data were obtained and in which settings the studied subjects lived. We found two important factors that potentially compromise external validity of the outcomes collected: the presence of a strong taxonomic bias, and an overrepresentation of results from studies in parrots kept in laboratories. The term “taxonomic bias” refers to differences in our knowledge of certain species and the degree to which they are the subject of scientific investigation across a wide variety of biological fields (113). The fact that genera like *Amazona, Melopsittacus*, and *Nymphicus* received more research attention compared to other groups such as *Cacatua, Myiopsitta*, or *Agapornis* clearly demonstrates a bias in the scientific literature. Several factors might have contributed to this discrepancy. For instance, parrots like budgerigars (*Melospittacus undulatus*) and cockatiels (*Nymphicus hollandicus*) are easily found, possess lower economic value, are easy to handle and tend to have a short generation interval as they become sexually mature before one year. All these characteristics, typical of commonly used animal models, make these species good candidates to conduct scientific studies under laboratory conditions. Although lovebirds (*Agapornis* spp.) possess similar characteristics, only one study on this genus met our inclusion criteria. Amazon parrots (*Amazona* spp.) do not possess any of these characteristics, yet were the most studied species, with the second-highest number of significant and feasible outcome measures. This is explained not by the widespread study of these species, but rather by the large amount of information gathered from one lab at the University of California, Davis, which published several studies on this taxon. We retrieved scarce data for the genera *Calyptorhynchus, Guaruba*, and *Loriculus*, but this is not surprising as these genera are mostly kept in zoos and breeding facilities and are rarely used in laboratory settings or kept as companion parrots. Nonetheless, no studies could be found for certain genera that are commonly kept in captivity, including *Psittacula, Pionites, Eolophus* and *Eclectus*.

Studies conducted on multiple genera included these taxa along with many others, however the information obtained from these studies should be evaluated carefully before being used for individual species assessments. Some taxonomic groups have in fact specific needs and show different sensitivities when exposed to similar environmental stimuli, making generalisation of findings to other genera sometimes difficult or irrelevant. It should be emphasized that data extrapolation should be performed with caution even within the same genus, as for example *Amazona, Ara* and *Cacatua* each contain several species adapted to different natural habitats and showing distinct behaviours within the same genus (114). As such, extrapolating findings to other species, even within the same genus, may be improper and counterproductive, emphasizing the need for further research to bridge these knowledge gaps for understudied species.

Our results show that certain welfare dimensions and categories were investigated only in a limited number of genera. For instance, we identified outcomes such as abnormal, locomotor, exploratory, and foraging behaviours for *Amazona, Ara*, and *Melopsittacus*, but these parameters were not described in *Agapornis, Cacatua*, and *Myiopsitta*. It is important to underline that the absence of information related on some taxa in our findings was not necessarily due to a lack of scientific studies but rather to a lack of significant results. For instance, we found that the provision of physical enrichment was linked to changes of preening in the genera *Amazona, Ara* and *Pyrrhura* (115-118), but not in *Nymphicus* (119, 120). In fact, none of the results obtained from this genus presented a significant p-value. This could mean that cockatiels, unlike other parrots’ species, do not show changes in preening behaviour in these situations or, alternatively, that experimental setups and methods applied in the studies were not able to detect such changes.

As mentioned above, more than half of the outcomes were found from studies conducted in laboratory settings. Laboratories are highly controlled environments where daily routines related to animal care and testing are highly standardized. Such settings theoretically allow to render results more reproducible, but only in the case that characteristics are similar to those from which original data were extrapolated. This condition, defined as “standardization fallacy”, can lead to a decrease of external validity (121). In our case, external validity is important as parrot welfare indicators should ideally be applicable across parrots of different species and leaving in various living conditions. Although zoos, shelters, rehabilitation centres, and breeding facilities offer settings that are not as standardized as in laboratories, they still differ greatly from domestic environments, which raises caution in extrapolating these findings to companion parrots.

#### Companion parrots

We found limited information related to companion parrots in terms of the number and variety of outcomes measures. In fact, the number of feasible and significant outcomes retrieved from companion parrots was less than 5% of the total outcome measures collected, and 75% of those were related to feather damaging behaviour. Moreover, half of these outcomes for companion parrots were obtained from only three genera: *Psittacus, Cacatua* and *Agapornis*, with *Psittacus* being the only genus that covered the entire breath of welfare categories, and outcomes for the other two genera being restricted to those related to feather damaging behaviour. While these three genera are among the most commonly kept companion parrot species, several other species, such as macaws, amazon parrots, conures, caiques, parrotlets, budgerigars, and other parakeets are also popular as companions, for which no significant and feasible outcomes were identified. An additional problematic aspect is that almost all outcome measures for companion parrots were obtained through questionnaires. Prospective, case-control studies might be challenging to perform with companion parrots under experimental circumstances as parrots may be harder to recruit, and possibly be less adaptable to new environments and/or unfamiliar humans compared to dogs and cats, which could lead to altered behavioural responses. Questionnaires may represent the best way to study this specific cohort and gather large amounts of data while preventing potential discomfort in the studied parrots. However, questionnaires are also highly sensitive to bias (122), and therefore results obtained using this method should be applied cautiously and possibly require complementary experimental testing. Overall, these results emphasize the need for more research on companion parrots, especially on species and welfare dimensions that have been underrepresented in this cohort.

#### Limitations

This systematic review presents some limitations. The first is a lack of terms used to create the search queries, especially those related to medical conditions. This would certainly allow to increase the final number of studies retrieved; however, due to the high number of diseases and pathologic conditions, we decided to select terms related to specific medical conditions. The second important limiting factor was the fact that data collection was carried out by only one person (AP), which might have led to missing information or to systematic errors that reduced the accuracy of the results presented. The third and last limitation is that we assessed internal validity only considering the p-value significance. A p-value in itself might be a necessary parameter, but it is insufficient to ascertain internal validity as it can be influenced by confounding factors, leading to false negative or false positive results. Further studies should include the effect size as a parameter to assess internal validity with higher confidence, but this was not possible in this study based on the information provided by authors.

## Conclusion

The purpose of this systematic review was to identify valid and feasible welfare indicators for companion parrots. The lack of information contained in the publications made it difficult to assess the internal validity of the outcome measures. We also noticed potentially low external validity due to taxonomic bias and an overrepresentation of studies on parrots kept in laboratories. These challenges to ascertain validity prevented us from establishing a definitive list of reliable and useful welfare indicators for companion parrots. Nevertheless, this systematic review helps to summarize the current state of scientific knowledge on aspects relevant to parrot welfare and identifies potential welfare indicators that require further validation through further research efforts.

## Acknowledgments

We would like to thank Marta Trogu and Lewis Urquhart for proofreading the manuscript, Suzanne Truong for proofreading and for offering valuable comments on figures and writing style, and Patrizia Piotti for providing advice and constructive feedback about assessment of internal validity.

## Author Contributions

Conceptualization, J.-L.R., A.P.; methodology, all authors; data collection, A.P.; statistical analysis, A.P; data curation, A.P.; writing—original draft preparation, A.P.; writing—review and editing A.P, Y.R.A.v.Z., J.-L. R.; supervision, Y.R.A.v.Z., J.-L. R.

