## Supplemental file for "What We (Don’t) Know about Parrot Welfare: A Systematic Literature Review"

#### Material and Methods

##### Search Query creation

A search query was created using the terms reported in table S1 and additional terms included in the thesaurus of the databases. Filters related to language, publication type and terms to exclude from the results of the systematic search (e.g., "wild birds") were also included in the search queries.

**Table S1**. Terms used to create the search queries.

| Search components | Terms |
|---|---|
| **Population** | parrot, psittacines, *Psittacids*, cockatoo, macaw, parakeet, budgerigar, cockatiel, *Ara*, *Cacatua*, *Psittacus*, african grey parrot, "grey parrot", amazon parrot, lovebirds, *Poicephalus, Agapornis*, *Psittaccula*, *Eclectus*, conure, caique. |
| **Intervention** | enrichment, environmental enrichment, social enrichment, nutritional enrichment, physical enrichment, sensory enrichment, deprivation, diet, nutrition, malnutrition, play, toy, puzzle, play activity, foraging, foraging toy, foraging activity, social activity, activity, stimulation, social bond, social relation, cognitive stimulation, attachment, hand rearing, parent rearing, co-parenting |
| **Outcomes** | feather picking, feather plucking, feather damaging, self-damaging, self-mutilation, pododermatitis, atherosclerosis, metabolic bone disease, body condition, obesity, nutritional deficiency, injuries, disease, stereotypes, stereotypical behaviours, abnormal behaviours, behavioural disorders, destructive behaviour, egg laying, reproduction, fertility, biting, screaming, excessive vocalization, natural behaviours, aggressive, emotion, emotional state, positive state, negative state, stress, distress, hormones, corticosterone, lifespan, life expectancy, longevity, aging |

For each database, we applied filters in order to exclude irrelevant studies and optimize the results of the systematic search. These filters included the language restriction (only English), the exclusion of studies focused on wild parrots and reviews or case studies. From a preliminary systematic search, we identified irrelevant studies (e.g. studies focused on bacteria, plants, mammals etc.), and therefore added online filters offered by the databases. This included filters related to the topic, authors name (e.g. Parrot et al) and studies category (e.g. Fishery, Marine Freshwater Biology, Sport Sciences etc.). In addition, on PubMed we applied relevant MeSH terms and the filter "Title/Abstract", on CAB direct relevant organism descriptors and topic terms.

##### Systematic Search Queries

Below are the search queries used on the 3 databases.

##### PubMed

(((((((((((((((((((((((((((((Parrot[Title/Abstract]) OR (Psittacines[Title/Abstract])) OR (Psittacids[Title/Abstract])) OR (Psittacidae[Title/Abstract])) OR (Psittacinae[Title/Abstract])) OR (Cacatuidae[Title/Abstract])) OR (Cockatoo[Title/Abstract])) OR (Macaw[Title/Abstract])) OR (Parakeet[Title/Abstract])) OR (Budgerigar[Title/Abstract])) OR (Cockatiel[Title/Abstract])) OR) OR (Cacatua[Title/Abstract])) OR (Psittacus[Title/Abstract])) OR (Grey parrot[Title/Abstract])) OR (Amazona[Title/Abstract])) OR (lovebird[Title/Abstract])) OR (Agapornis[Title/Abstract]) )) OR (Psittacula[Title/Abstract])) OR (Conure[Title/Abstract])) OR (Caique[Title/Abstract])) OR (Ecletus[Title/Abstract])*) OR (Poichephalus[Title/Abstract])*) OR ("Parrots"[MAJR])) OR ("Cockatoos"[MAJR])) AND (((((((((((((((((((((((((((((((((((((((((((((((((((((((((((((((((((("Feather picking") OR ("feather plucking")) OR ("feather damaging")) OR ("feather destruction")) OR (self-damaging)) OR (self-mutilation)) OR (automutilation)) OR (pododermatitis)) OR (atherosclerosis)) OR ("metabolic bone desease")*) OR ("Body condition")) OR (obesity)) OR (injury)) OR (disease)) OR ("nutritional deficiency")) OR (Stereotypes)) OR ("stereotypical behaviour"))

OR ("abnormal behaviour")) OR ("repetitive behaviour")*) OR ("behavioural disorder")) OR ("destructive behaviour")) OR ("natural behaviour")) OR ("egg laying")) OR (reproduction)) OR (fertility)) OR (aggression)) OR (emotion)) OR ("emotional state")) OR (biting)) OR (screaming)) OR ("excessive vocalization")) OR ("positive state")) OR ("negative state")) OR (Stress)) OR (distress)) OR (hormones)) OR (corticosterone)) OR (Lifespan)) OR ("life expectancy")) OR (longevity)) OR (aging)) OR ("Bird Diseases/diagnosis"[MeSH])) OR ("Cognition"[MeSH])) OR ("Emotions"[MAJR])) OR ("Parrots/physiology"[MAJR])) OR ("Vocalization, Animal"[MAJR])) OR ("Feathers/injuries"[MAJR])) OR ("Corticosterone/blood"[MeSH])) OR ("Diagnosis, Differential"[MeSH])) OR ("Aggression/physiology"[MAJR])) OR ("Stereotyped Behavior"[MAJR])) OR ("Impulsive Behavior"[MeSH])) OR ("Behavior, Animal"[MeSH])) OR ("Feeding Behavior/physiology"[MAJR])) OR ("Amazona/metabolism"[MAJR])) OR ("Amazona/physiology"[MAJR])) OR ("Amazona/growth and development"[MAJR])) OR ("Animal Feed/analysis"[MAJR]))) OR ((((((((((((((((((((((((Parrot[Title/Abstract]) OR (Psittacines[Title/Abstract])) OR (Psittacids[Title/Abstract])) OR (Psittacidae[Title/Abstract])) OR (Psittacinae[Title/Abstract])) OR (Cacatuidae[Title/Abstract])) OR (Cockatoo[Title/Abstract])) OR (Macaw[Title/Abstract])) OR (Parakeet[Title/Abstract])) OR (Budgerigar[Title/Abstract])) OR (Cockatiel[Title/Abstract])) OR) OR (Cacatua[Title/Abstract])) OR (Psittacus[Title/Abstract])) OR (Grey parrot[Title/Abstract])) OR (Amazona[Title/Abstract])) OR (lovebird[Title/Abstract])) OR (Agapornis[Title/Abstract]) )) OR (Psittacula[Title/Abstract])) OR (Conure[Title/Abstract])) OR (Caique[Title/Abstract])) OR (Ecletus[Title/Abstract])*) OR (Poichephalus[Title/Abstract])*) OR ("Parrots"[MAJR])) OR ("Cockatoos"[MAJR])) AND ((((((((((((((((((((((((((((((((Enrichment) OR ("Environmental Enrichment")) OR ("Social Enrichment")) OR ("Nutritional Enrichment")) OR ("Physical Enrichment")) OR ("sensory enrichment")) OR (Deprivation)) OR (Diet)) OR (Nutrition)) OR (Malnutrition)) OR (Play)) OR (Toy)) OR (Puzzle)) OR ("Play activity")) OR (Foraging)) OR ("Foraging toy")) OR ("foraging activity")) OR ("social activity")) OR (activity)) OR (stimulation)) OR ("social bond")) OR ("social relation")) OR ("cognitive stimulation")) OR (attachment)) OR ("Hand rearing")) OR ("parent rearing")) OR (co-parenting)) OR ("Behavior, Animal/drug effects"[MeSH])) OR ("Animal Nutritional Physiological Phenomena/physiology"[MeSH])) OR ("Acoustic Stimulation"[MeSH])) OR ("Social Isolation/psychology"[MAJR])) OR ("Play and Playthings"[MAJR]))) AND (english[Filter] OR french[Filter] OR italian[Filter])) NOT (Review[Publication Type])) NOT (Case Reports[Publication Type])) NOT ("wild birds") AND (english[Filter] OR french[Filter] OR italian[Filter])

*Misspelled terms

#### CAB direct

(((((((((up:("Cacatuidae" or "Psittacidae")) OR ((up:("Psittaciformes" or "Psittacus")) OR (((((((Parrot) OR ( Psittacines) OR ( Psittacids) OR (Psittacidae) OR (Psittacinae) OR (Cacatuidae) OR ( Cockatoo) OR (Macaw) OR (Parakeet) OR (Budgerigar) OR (Cockatiel) OR (Ara) OR (Cacatua) OR (Psittacus) OR (Grey parrot) OR ( lovebird) OR ( Agapornis) OR (Psittacula) OR (Conure) OR (Caique) OR (Ecletus)* OR (Poichephalus)*))) OR (od:("Ara" or "Amazona" or "Cacatua" or "Eclectus roratus" or "Nymphicus hollandicus" or "Psittacula" or "Myiopsitta monachus" or "Cacatuidae" or "Nymphicus" or "Pionites" or "Psittacus" or "Eclectus" or "Aratinga")))) OR (od:("Cacatua" or "Agapornis" or "budgerigars"))))) AND ((de:("behaviour problems")) OR (((id:("chewing")) OR ((((((("Feather picking") OR ("feather plucking") OR ( "feather damaging") OR ("feather destruction") OR (self-damaging) OR (self-mutilation) OR ("automutilation") OR (pododermatitis) OR (atherosclerosis) OR ("Body condition") OR (obesity) OR (injury) OR (disease) OR ("nutritional deficiency") OR ( Stereotypes) OR ( "stereotypical behaviour") OR ("abnormal behaviour") OR ("repetetive behaviour")* OR ("behavioural disorder") OR ("destructive behaviour") OR ( "natural behaviour") OR ("egg laying") OR (reproduction) OR (fertility) OR (aggression) OR (emotion) OR ("emotional state") OR (biting) OR (screaming) OR ("excessive vocalization") OR ("positive state") OR ("negative state") OR (Stress) OR (distress) OR (hormones) OR (corticosterone) OR (Lifespan) OR ("life expectancy") OR (longevity) OR (aging))) OR (de:("corticosterone" or "behaviour" or "feather pecking" or "animal behaviour" or "egg production")))) OR (id:("pterotillomania" or "cage birds" or "feather damaging behaviour" or "captive animals")))) OR (id:("deviant behaviour" or "abnormal behavior")))) OR (id:("chewing"))))) OR (((((((Enrichment) OR ( "Environmental Enrichment") OR ("Social Enrichment") OR ("Nutritional Enrichment") OR ("Physical Enrichment") OR ( "sensory enrichment") OR (Deprivation) OR (Diet) OR (Nutrition) OR (Malnutrition) OR (Play) OR (Toy) OR (Puzzle) OR (Play activity) OR (Foraging) OR (Foraging toy) OR (foraging activity) OR (social) OR (activity) OR (stimulation) OR (social bond) OR ( social relation) OR (cognitive stimulation) OR (attachment) OR (Hand rearing) OR (parent rearing) OR (co-parenting))) OR (de:("enrichment" and "physical activity" and "quality of life" and "ornamental birds" and "toys" and "animal welfare" and "foraging")))) OR (de:("exercise" or "animal nutrition" or "diets" or "ontogeny"))) AND ((up:("Cacatuidae" or "Psittacidae")) OR ((up:("Psittaciformes" or "Psittacus")) OR (((((((Parrot) OR ( Psittacines) OR ( Psittacids) OR (Psittacidae) OR (Psittacinae) OR (Cacatuidae) OR ( Cockatoo) OR (Macaw) OR (Parakeet) OR (Budgerigar) OR (Cockatiel) OR (Ara) OR (Cacatua) OR (Psittacus) OR (Grey parrot) OR (Amazona) OR ( lovebird) OR ( Agapornis) OR (Psittacula) OR (Conure) OR (Caique) OR (Ecletus)* OR (Poichephalus)*))) OR (od:("Ara" or "Amazona" or "Cacatua" or "Eclectus roratus" or "Nymphicus hollandicus" or "Psittacula" or "Myiopsitta monachus" or "Cacatuidae" or "Nymphicus" or "Pionites" or "Psittacus" or "Eclectus" or "Aratinga")))) OR (od:("Cacatua" or "Agapornis" or "budgerigars")))))) NOT (Review)) NOT (de:("wild birds"))) NOT ("Case Study")) NOT (de:("DNA cloning" or "interferon-gamma"))) NOT (de:("guided tours" or "cultural tourism" or "ecotourism"))) AND ( ((NOT (organism-descriptor:(( "Protozoa" OR "mice" OR "fowls" OR "fishes" OR "plants" ) ))) (NOT (topic:(( "aquatic organisms" OR "aquatic species" OR "aquatic animals" OR "case reports" OR "forests" OR "bacterium" ) ))) (NOT (broader-term:(( "mammals" OR "Spermatophyta" OR "angiosperms" OR "plants" ) ))) ))) AND ( ((topic:(( "cage birds" OR "aviary birds" OR "behavior" OR "behaviour" OR "animal behavior" OR "pet animals" OR "pets" OR "animal behaviour" ) )) (NOT (topic:(( "Viral diseases" OR "viral infections" ) ))) (language:(( "English" OR "French" OR "Italian" ) ))))

*Misspelled terms

#### Web of Science

Web of Science did not allow to create a search query containing more than 100 terms, for this reason terms related to the intervention and the outcomes were reduced.

((((((((((((((((((((ALL=(Parrot)) OR ALL=(Psittacines)) OR ALL=(Psittacids)) OR ALL=(Psittacidae)) OR ALL=(Psittacinae)) OR ALL=(Cacatuidae)) OR ALL=(Cockatoo)) OR ALL=(Macaw)) OR ALL=(Parakeet)) OR ALL=(Budgerigar)) OR ALL=(Cockatiel)) OR ALL=(Ara)) OR ALL=(Cacatua)) OR ALL=(Psittacus)) OR ALL=("Grey parrot")) OR ALL=(Amazona)) OR ALL=(lovebird)) OR ALL=(Agapornis)) OR ALL=(Psittacula)) OR ALL=(Conure)) OR ALL=( Caique)) OR ALL=(Ecletus)*) OR ALL=(Poichephalus)* AND (((((((((((((((((ALL=(Enrichment)) OR ALL=("sensory enrichment" )) OR ALL=(Deprivation)) OR ALL=(Diet)) OR ALL=(Nutrition)) OR ALL=(Malnutrition)) OR ALL=(Play)) OR ALL=( Toy)) OR ALL=(Puzzle)) OR ALL=(Foraging)) OR ALL=( "social")) OR ALL=(activity)) OR ALL=(stimulation)) OR ALL=("social bond")) OR ALL=("cognitive stimulation")) OR ALL=("attachment")) OR ALL=(rearing)) OR ALL=( co-parenting) AND (((((((((((((((((((((((((((((((((ALL=(Feather)) OR ALL=(damage)) OR ALL=(destruction)) OR ALL=(mutilation)) OR ALL=(pododermatitis)) OR ALL=(atherosclerosis)) OR ALL=( "metabolic bone desease")*) OR ALL=("Body condition")) OR ALL=(obesity)) OR ALL=( injury)) OR ALL=( disease)) OR ALL=( "nutritional deficiency")) OR ALL=(Stereotypes)) OR ALL=("stereotypical behaviour")) OR ALL=("abnormal behaviour")) OR ALL=("repetitive behaviour")*) OR ALL=( "behavioural disorder")) OR ALL=("natural behaviour")) OR ALL=("egg laying")) OR ALL=("reproduction")) OR ALL=( fertility)) OR ALL=(aggression)) OR ALL=("emotional state")) OR ALL=(biting)) OR ALL=(screaming)) OR ALL=( "excessive vocalization")) OR ALL=( "positive state")) OR ALL=("negative state")) OR ALL=( Stress)) OR ALL=( distress)) OR ALL=( hormones)) OR ALL=(corticosterone)) OR ALL=(Lifespan)) OR ALL=( "life expectancy")) OR ALL=(longevity)) OR ALL=(aging)

*Misspelled terms

#### Inclusion and Exclusion Criteria

To select articles that fit with the research questions, we applied the eligibility criteria described in table S2.

**Table S2.** Inclusion and exclusion criteria used to screen the articles by title/abstract reading and full-text reading.

| | | |
|---|---|---|
| **Population** | Inclusion criteria | Species: all species belonging to the order Psittaciformes |
| | | Parrots living as companion animals or laboratory animals, in zoos, shelters or breeding centres |
| | | Demographic factors: all ages, both sexes |
| | Exclusion criteria | Wild parrots |
| **Intervention** | Inclusion criteria | Enrichment: enriched parrot vs not enriched (between or within subject designs)<br>Social enrichment (e.g. intra-interspecific interactions, hand-raised vs parent raise), physical enrichment (e.g. foraging toys, changes in the aviary/cage/room), cognitive enrichment (problem solving activities, training etc.), nutritional enrichment (vegetable and/or fruits as well as seeds), etc. |
| | | Vet Treatments: treated vs not treated |
| | | Examination: In Vivo, Post-mortem |
| | | Diet manipulation: increased/decreased levels of cholesterol/fibres/fat etc. |
| | | Questionnaires |
| | | Behavioural tests: novel object, open field, flight training, etc. |
| | | Personality tests: only when result are related to the outcomes described below |
| **Outcomes measures** | Inclusion criteria | Problematic behaviours: aggressiveness (e.g. bites), fear-related behaviour (avoidance behaviour), abnormal behaviours (self-mutilation, feather plucking, stereotypies, incessant screaming). |
| | | Activity level: resting, exploratory behaviour, foraging behaviour |
| | | Social behaviours: play, allopreening, interspecific interactions, human interactions |
| | | Vocal behaviours |
| | | "Body language": feather ruffling, body posture |
| | | Health measures: body weight, plumage quality, presence of diseases, mortality |
| | | Behaviour related to basic needs: eating, drinking, resting |
| | | Chick mortality only when related to aggressive interactions |
| | Exclusion criteria | Reproductive parameters: egg hatchability, num. of eggs laid |
| | | Chicks-related: development parameters and nutritional requirements |
| **General** | Inclusion criteria | Only English |
| | | Publication date restriction: none |
| | Exclusion criteria | Methodological details: studies that lacked statistical analysis |
| | | Publication type: reviews, conference abstracts, book chapters |
| | | Irretrievable studies |
| | | Study design: case studies, studies with less than 5 subjects |

#### Full-text screening

Two different reviewers (AP and J-LR) screened the 140 articles. The reviewers, using the exclusion and the exclusion criteria, independently select the papers that considered eligible. After the full-text screening, the reviewers compared their two lists of eligible studies and decide which ones to exclude or include. In case of disagreement between the reviewers a study was considered not eligible.

### Data collection

#### Descriptions of the validity measures

**Table S2**. Definitions of the validity parameters used to assess the risk of bias of outcome measures.

| Variable | Description | **Levels** |
|---|---|---|
| **Inter-observer reliability** | Were the outcomes tested for inter-reliability? | **Yes**: authors reported in the main text that the outcome was tested for inter-observer reliability<br>**No**: authors reported in the main text that the outcome was not tested for inter-observer reliability<br>**Unspecified**: authors did not report the information in the main text<br>**Not Possible**: the experimental set up did not allow to test for inter-observer reliability |
| **Intra-observer reliability** | Were the outcomes tested for intra-reliability? | **Yes**: authors reported in the main text that the outcome was tested for intra-observer reliability<br>**No**: authors reported in the main text that the outcome was not tested for intra-observer reliability<br>**Unspecified**: authors did not report the information in the main text<br>**Not Possible**: the experimental set up did not allow to test for intra-observer reliability |
| **P-value significance** | Was the p-value significant? | **Significant**: $p < 0.05$<br>**Not significant**: $p > 0.05$<br>Cases when the p-value significance were unclear were considered not significant |
| **Random group assignment** | Were the subjects randomly assigned to experimental and control groups? | **Yes**: authors reported random group assignments of the subjects in the main text<br>**No**: authors reported in the main text that the subjects were not randomly assigned to groups<br>**Unspecified**: authors did not report the information in the main text<br>**Not Possible**: the experimental set up did not allow to randomly assign subjects to groups |
| **Condition balancing** | Were conditions balanced between subjects and/or groups? | **Yes**: authors reported in the main text that experimental conditions were balanced between subjects and/or groups<br>**No**: authors reported in the main text that experimental conditions were not balanced between subjects and/or groups<br>**Unspecified**: authors did not report the information in the main text<br>**Not Possible**: the experimental set up did not allow to balance conditions between subjects and/or groups |
| **Observer blindness** | Was the outcome assessor blinded during data collection? | **Yes**: authors reported in the main text that the assessor was blinded<br>**No**: authors reported in the main text that the assessor was not blinded<br>**Unspecified**: authors did not report the information in the main text<br>**Not Possible**: the experimental set up did not allow to blind the assessor |

#### Welfare dimensions and outcome categories

**Table S3**. Welfare dimensions and corresponding outcomes categories

| Welfare Dimensions | Outcomes Categories |
|---|---|
| **Body measurements** | Indirect measures of feather damaging behaviours, body condition, feather colour |
| **Physiological parameters** | Stress-related, metabolic, vitamin D-related, lipid-related, immune system-related, body temperatures, others |
| **Abnormal and fear-related behaviour** | Feather-damaging behaviours, fear-related, stereotypies, incessant screaming |
| **Maintenance behaviour** | Feeding, drinking, resting, self-care |
| **Locomotor behaviour** | Fly, position occupied in the cage, locomotion, inactivity |
| **Exploratory and foraging behaviour** | Cognitive stimulation, enrichment interaction, foraging, environment/object preference, reaction to new environment, reaction to novel objects |
| **Diseases and pathologic conditions** | - |
| **Social behaviour** | Allopreening, aggressive behaviours, human-animal interaction, facial and body displays, sexual behaviours, social dynamics, vocalizations |

#### Subjects' living conditions

**Table S4**. Description of the living conditions of the subjects tested in the eligible studies.

| **Living condition** | **Definition** |
|---|---|
| Breeding facility | Parrots living in private or public breeding facilities |
| Companion animals | Parrots that live with humans or within human social structures where they are provided with some, or all, of their needs. They are considered to play a primarily social role within the household or community (1) |
| Laboratory animals | Parrot kept for research purposes at universities or laboratories |
| Multiple | Studies focused on parrots kept in multiple living conditions |
| Rehabilitation centre | Parrots kept in rehabilitation centres |
| Shelter | Parrots, previously kept as companion, that lives in shelters |
| Zoo | Parrots living in zoo with public or private aviaries |

### Results

**Table S5.** Genera and corresponding number of studies grouped according to the animals' living conditions.

| **Living condition** | **Number of Studies** | **Genera (number of studies)** |
|---|---|---|
| Breeding facility | 6 | *Amazona* (1), *Ara* (1), *Nymphicus* (3), *Guaruba* (1) |
| Companion animals | 21 | *Agapornis (1), Cacatua* (2), Multiple (11), Nymphicus (1), *Psittacus* (7) |
| Laboratory-kept animals | 48 | *Amazona* (22), *Melopsittacus* (14), Multiple (1), *Myiopsitta* (2), *Nymphicus* (7), *Platycercus* (1), *Psittacus* (1) |
| Multiple | 1 | *Amazona* (1) |
| Rehabilitation centre | 3 | *Amazona* (3) |
| Shelter | 2 | *Psittacus* (2) |
| Unknown | 7 | *Amazona* (1), *Ara* (1), *Myiopsitta* (1), Multiple (2), *Loriculus* (1), *Psittacus* (1) |
| Zoo | 10 | *Amazona* (1), *Ara* (4), *Cacatua* (1), *Calyptorhynchus* (1), Multiple (2), *Pyrrhura* (1) |

#### Genera and living conditions represented in the studies

Eligible studies covered 13 genera, of which 10 belonged to the superfamily *Psittaccoidea* and 3 to the superfamily *Cacatuoidea.* None of the studies investigated the welfare of species belonging to the superfamily *Strigopoidea*. *Amazona* was the genus most investigated, representing 29 studies (29.6%), followed by the genera *Melopsittacus*, *Nymphicus*, and *Psittacus* with 14 (14.3%), 11 (11.2%), and 10 studies (10.2%) respectively (Figure S1, Table S8). A lesser number of studies focused on the genera *Ara* (n=6; 6.1%), *Cacatua* (n=3; 3.1%), and *Myiopsitta* (n=3; 3.1%), while six other genera were represented by only one study each (Figure S1, Table S8). Sixteen studies (16.3%) investigated parrot welfare in multiple genera, including those as previously mentioned, as well as the *Eclectus* (6 studies) and *Poicephalus* (6 studies) genera (Figure S1, Table S7 for a full list of species included in the category "multiple", Table S8).

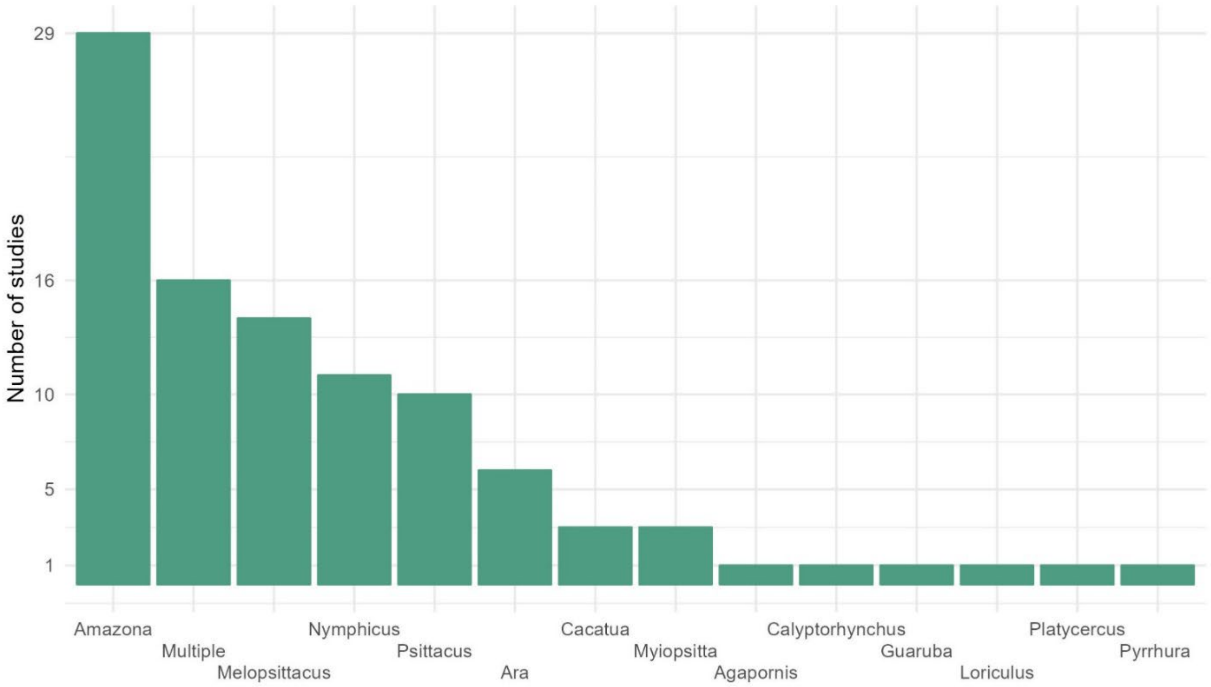


**Figure S1.** Number of studies grouped by parrots' genera. Barplot showing on the y-axis the number of studies for each parrot's genera.

**Table S6.** Parrots' genera investigated in studies focused on multiple species, followed by the number of studies that reported them.

| Genus | Number of Studies | Genus | Number of Studies | Genus | Number of Studies | Genus | Number of Studies |
|---|---|---|---|---|---|---|---|
| *Agapornis* | 6 | *Cyanoramphus* | 3 | *Nadayus* | 1 | *Psittacara* | 1 |
| *Alisterus* | 1 | *Deroptyus* | 1 | *Neophema* | 2 | *Psittacula* | 4 |
| *Amazona* | 10 | *Diopsittaca* | 2 | *Neopsittacus* | 1 | *Psittaculorostris* | 1 |
| *Anodorhynchus* | 2 | *Eclectus* | 6 | *Northiella* | 1 | *Psittacus* | 10 |
| *Aprosmictus* | 1 | *Enicognathus* | 1 | *Nymphicus* | 7 | *Psitteuteles* | 2 |
| *Ara* | 8 | *Eolophus* | 2 | *Opopsitta* | 1 | *Psittaculorostris* | 1 |
| *Aratinga* | 3 | *Eos* | 2 | *Orthopsittaca* | 1 | *Psittinus* | 1 |
| *Barnadius* | 2 | *Eupsittula* | 1 | Other[1] | 1 | *Purpureicephalus* | 1 |
| *Bolborhynchus* | 3 | *Forpus* | 4 | Parakeet[2] | 2 | *Pyrrhura* | 2 |
| *Brotogeris* | 1 | *Glossopsitta* | 1 | *Pionites* | 3 | *Tanygnathus* | 2 |
| *Cacatua* | 6 | *Guaruba* | 1 | *Pionopsitta* | 1 | *Trichoglossus* | 3 |
| *Callocephalon* | 1 | *Lathamus* | 1 | *Pionus* | 3 | *Triclaria* | 1 |
| *Chalcopsitta* | 2 | *Lorie* | 1 | *Platycercus* | 3 | Undetermined[1] | 2 |
| *Charmosyna* | 1 | *Lorius* | 2 | *Poicephalus* | 7 | Unknown[1] | 1 |
| Cockatoo[2] | 3 | Macaw[2] | 2 | *Primolius* | 1 | | |
| Conure[2] | 3 | *Melopsittacus* | 4 | *Probosciger* | 1 | | |
| *Coracopsis* | 1 | *Myiopsitta* | 4 | *Psephotus* | 2 | | |
| *Cyanoliseus* | 2 | *Neopsephotus* | 1 | *Pseudeos* | 2 | | |

[1] *Description reported by the authors.*
[2] *Authors reported common name instead of scientific name.*

**Table S7**. Number of studies and number of outcomes for each genus.

| ***Genera*** | **Studies (%)** | **Outcomes (%)** | **Significant and feasible outcomes (%)** | ***Outcomes (%); only companion*** | ***Significant and feasible outcomes (%); only companion*** |
|---|---|---|---|---|---|
| | **Total= 98** | **Total= 1512** | **Total= 572** | **Total= 340** | **Total= 68** |
| *Agapornis* | 1 (1.2%) | 21 (1.39%) | 3 (0.52%) | 21 (6.18%) | 3 (4.41%) |
| *Amazona* | 29 (29.59%) | 320 (21.16%) | 128 (22.34%) | 12 (3.53%) | / |
| *Ara* | 6 (6.12%) | 142 (9.39%) | 72 (12.57%) | / | / |
| *Cacatua* | 3 (3.06%) | 69 (4.56%) | 12 (2.09%) | 65 (19.12%) | 9 (13.24%) |
| *Calyptorhynchus* | 1 (1.02%) | 6 (0.4%) | 4 (0.7%) | / | / |
| *Guaruba* | 1 (1.02%) | 3 (0.2%) | 1 (0.17%) | / | / |
| *Loriculus* | 1 (1.02%) | 3 (0.2%) | 3 (0.52%) | / | / |
| *Melopsittacus* | 14 (14.28%) | 287 (20.63%) | 150 (26.22%) | / | / |
| *Multiple* | 16 (16.32%) | 287 (18.98%) | 54 (9.42%) | 150 (44.12%) | 32 (47.06%) |
| *Myiopsitta* | 3 (3.06%) | 48 (3.17%) | 9 (1.57%) | / | / |
| *Nymphicus* | 11 (11.22%) | 148 (9.79%) | 65 (11.34%) | 4 (1.18%) | / |
| *Platycercus* | 1 (1.02%) | 26 (1.72%) | / | / | / |
| *Psittacus* | 10 (10.20%) | 133 (8.80%) | 54 (9.44%) | 88 (25.88%) | 24 (35.29%) |
| *Pyrrhura* | 1 (1.02%) | 19 (1.26%) | 17 (2.97%) | / | / |

#### Welfare dimensions represented in the studies

The most common welfare dimensions investigated across all studies, in decreasing order, were "body measurements" (35 studies), followed by "social behaviours" (34 studies), "physiological parameters" (31 studies), "maintenance behaviours" (29 studies), "exploratory and foraging behaviours" (28 studies), "locomotor behaviours" (20 studies), "abnormal and fear-related behaviours" (16 studies), and "diseases and pathologic conditions" (11 studies) (Figure S2 and Table S9).

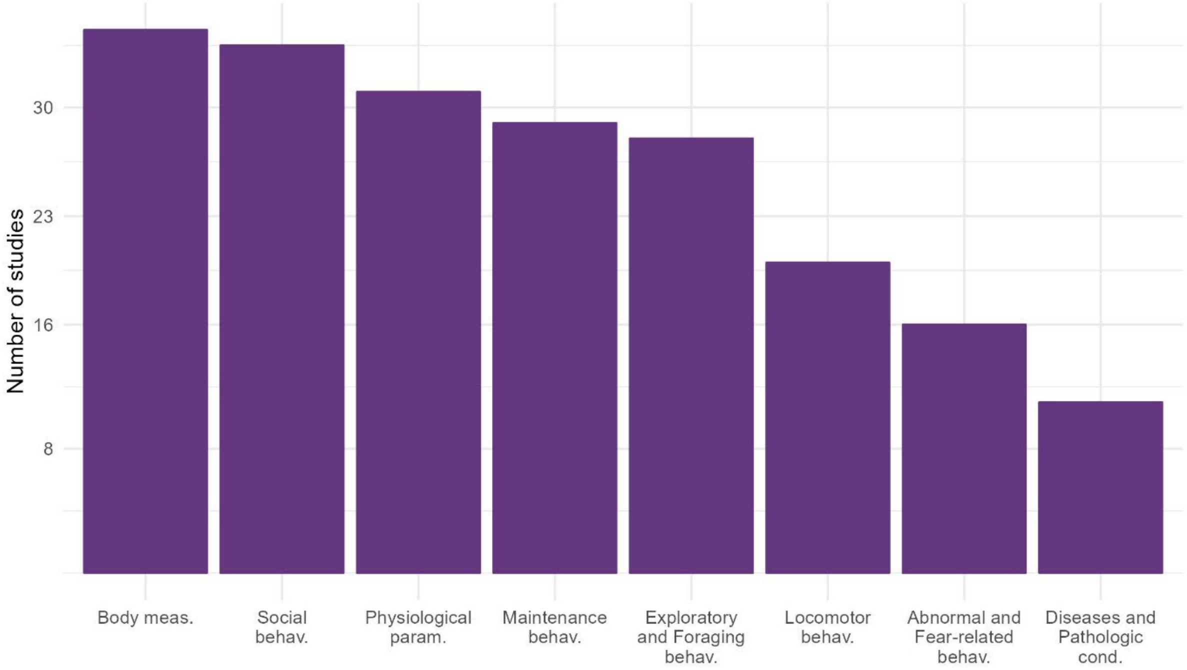


**Figure S2.** Studies grouped by welfare dimensions. Barplot showing on the y-axis the number of studies for each welfare dimensions. Behav.=behaviour, meas.=measurements, param.=parameters, cond.=conditions.

**Table S8.** Number of studies and outcomes for each welfare dimension.

| *Welfare Dimensions* | **Studies (%)** | **Outcomes (%)** | **Significant and feasible outcomes (%)** | ***Outcomes (%); only companion*** | ***Significant and feasible outcomes (%); only companion*** |
|---|---|---|---|---|---|
| | | **Total= 1512** | **Total= 572** | **Total= 340** | **Total= 68** |
| *Abnormal and fear-related behaviours* | 16 (7.84%) | 160 (10.58%) | 88 (15.38%) | 34 (10%) | 8 (11.76%) |
| *Locomotor behaviours* | 20 (9.80%) | 149 (9.85%) | 83 (14.51%) | / | / |
| *Exploratory and foraging behaviours* | 28 (13.72%) | 212 (14.02%) | 93 (16.26%) | 1 (0.29%) | 1 (1.47%) |
| *Diseases and pathologic conditions* | 11 (5.39%) | 61 (4.03%) | / | 17 (5%) | / |
| *Maintenance behaviours* | 29 (14.21%) | 185 (12.24%) | 87 (15.21%) | 6 (1.76%) | 2 (2.94%) |
| *Body measurements* | 35 (17.15%) | 316 (20.90%) | 80 (13.99%) | 231 (67.94%) | 51 (75%) |
| *Physiological parameters* | 31 (15.19%) | 192 (12.70%) | / | 45 (13.24%) | / |
| *Social behaviours* | 34 (16.66%) | 237 (15.67%) | 141 (24.65%) | 6 (1.76%) | 6 (8.82%) |

#### Welfare categories represented in the studies

The outcome measures were grouped in 35 different welfare categories (Table S4). Of these, “indirect measures of feather-damaging behaviour” and “feeding” were covered by the highest number of studies (n=19), followed by “self-care” (n=18) and “stereotypies” (n=16) (Table S10). Most other categories were covered by 2 to 9 studies. Of all categories, “body surface temperature” was the least studied with only one study (Table S10).

**Table S10.** Significant and feasible outcome measures grouped by the welfare dimensions and by the 35 welfare categories. The table reports the number of outcomes for all welfare categories, the types of outcomes collected, and all genera investigated with their corresponding studies. The outcomes' names are kept as the authors reported them. D= duration, L= latency, F= frequency, P= Percentage, PP= proportion of parrots, U= unknown type of measurement.

| Welfare dimensions | Welfare Categories | Number of Outcomes | Types of outcomes | Feasible risk factors | Genera | Studies |
|---|---|---|---|---|---|---|
| **Abnormal and fear-related behaviours** | Fear-related | 3 (0.2%) | Phobic behaviours (U) | Cage size, perches' height, wild caught birds | *Psittacus* | (2) |
| | Incessant screaming | 8 (1.40%) | Incessant screaming (D, F) | Lack of social enrichment, cage size | *Melopsittacus* | (3, 4) |
| | Stereotypies | 77 (13.44%) | Oral stereotypies (P), total stereotypies (F, D, P), whole body stereotypies, self-biting, sham/wire chewing (F, D), beak grinding (F), biting (F), pacing (D, F, PP), route tracing (D, F, PP), spot pecking (D, F) | Social isolation, <5 weeks when removed from the nest, brain volume (species), cage size, lack of enrichment, lack of social interactions, bird sex, number of birds, unbalanced diet, rearing method, perches' material | *Amazona*, *Ara*, *Calyptorhynchus*, *Melopsittacus*, "Multiple", *Psittacus* | (2-14) |
| **Exploratory and foraging behaviours** | Cognitive | 6 (1.05%) | Proactive response towards novel objects, latency to reward during attention bias test, problem solving skills, responsiveness during discrimination task | Feather damaging behaviours, presence of unfamiliar humans, personality | *Amazona*, *Melopsittacus*, *Psittacus* | (15-17) |
| | Enrichment interaction | 24 (4.19%) | Environment interaction (F, P, PP), object destruction, visit enriched area (F, D), enrichment interaction (F, D, P) | Lack of offered choices, social isolation, bird sex, lack of enrichments, rearing method, inappropriate handling, personality | *Amazona*, *Ara*, "Multiple", *Nymphicus* | (5, 6, 9, 10, 14, 18-22) |
| | Foraging | 16 (2.79%) | Foraging (F, D, P), interaction with food (D), contrafreeloading | Lack of enrichment, personality | *Ara*, *Calyptorhynchus*, *Psittacus* | (12, 14, 23-25) |
| | Preference | 18 (3.14%) | Choice for pellet size or objects colour/size/material, perch diameter/position | Lack of offered choices | *Amazona*, *Loriculus* | (26-29) |
| | Reaction to new environment | 7 (1.22%) | Exploration (D, L), pattern of exploration, total distance/amount squares covered, number of visits of new environments, explorative tendency PCA axis* | Negative correlation with neophobic behaviours, feather-damaging behaviours, social isolation | *Melopsittacus*, *Psittacus* | (15, 17, 30) |
| | Reaction to novel objects | 22 (3.84%) | Novel object interaction (D, L), latency to feed in presence of novel object, latency to touch novel object | Rearing method, lack of enrichment (physical + foraging), social isolation, personality, species, dominance | *Amazona*, "Multiple" | (5, 31-33) |
| **Locomotor behaviours** | Flying | 26 (4.54%) | Flying (D, F, L, score), escape flight (D) | Cage size, social isolation, presence of feather damaging behaviours, lack of enrichments, bird sex, rearing method, insufficient physical activity, wing load (indirect measure of fat mass) | *Amazona*, *Melopsittacus*, "Multiple", *Pyrrhura* | (5, 9, 11, 30, 34-37) |
| | Inactivity | 9 (1.57%) | Inactivity (PP), stationary position (D) | Lack of enrichment, time of day, correlated with feather damaging behaviours | *Ara*, *Pyrrhura* | (19, 34) |
| | Locomotion | 34 (5.93%) | Locomotion (F, D), walking (F, D), climbing (D), general activity (D), hopping (D), number of area changes | Lack of social interactions, cage size, lack of enrichment, social isolation, inappropriate handling | *Amazona*, *Ara*, *Melopsittacus*, *Nymphicus*, *Pyrrhura*, | (4, 5, 8, 10, 11, 14, 19, 20, 22, 34, 38) |
| | Position occupied in the cage | 14 (2.44%) | Standing at the bottom of the cage (F), flights from the perch/wall to the ground (F, D), aggregated flights to the ground (D), fly between walls/perches (F), time spent on the ground, time spent 1m to 2m high, standing at the grid ceiling (F) | Elevated ambient temperature, lack of enrichment, lack of social interaction, feeder position in the cage | *Ara*, *Melopsittacus*, *Nymphicus*, *Pyrrhura*, | (10, 22, 34, 35, 39) |
| **Maintenance behaviours** | Drinking | 2 (0.35%) | Water intake | Unbalanced diet, Artificial light at night | *Amazona*, *Melopsittacus* | (40, 41) |
| | Feeding | 31 (5.41%) | Food consumption (D), food intake, time spent feeding per day (P), feeding (D, F), visit to feeding dish (F), ratio consumption (grams per bird) | Social isolation, elevated temperatures, food preference, lack of enrichment (physical/foraging), personality (vigilance), time of day, inappropriate handling, cage size, bird sex, courtship feeding, flight activity (frequency), lack of social interactions, unbalanced diet, artificial light at night | *Amazona*, *Ara*, *Melopsittacus*, *Myiopsitta*, Nymphicus, *Psittacus* | (11, 19-21, 24, 30, 35, 39-46) |
| | Self-care | 42 (7.33%) | Preening (D, L, P), bathing (D, L), tail wagging (F), puffing up the feathers (F), scratching (PP), cleaning the beak (PP), wing stretch (F) | Reduced flight ability, rearing method, lack of bathing opportunities, feather damaging behaviours, lack of enrichment, cage size, bird sex, time of day, lack of social interactions, social isolation | *Amazona*, *Ara*, *Melopsittacus*, "Multiple", *Psittacus*, *Pyrrhura* | (2-5, 8-11, 19, 21, 30, 34, 42) |
| | Resting | 12 (2.09%) | Resting/sleeping (D, F, P) | Lack of enrichment, cage size, lack of social interactions, inappropriate handling, time of day, social isolation | *Amazona*, *Ara*, *Melopsittacus*, *Nymphicus* | (4, 5, 14, 20, 21, 42, 44) |
| **Body measurements** | Body condition | 12 (2.09%) | Body mass, body weight, chest girth | Artificial light at night, lack of exercise, unbalanced diet, flight activity, feeding duration, wing load (indirect measure of fat mass), bird sex, general activity | *Amazona*, *Ara*, *Melopsittacus* | (35, 41, 46-48) |
| | Indirect measures of feather damaging behaviour | 66 (12.04%) | Presence/absence of feather damages, plumage score | Diet needing extensive handling, number of vacations per year, age, cage size, absence of foraging/chewable devices, lack of command training, personality, distance of the cage from the door, variety of the diet, heritability, hours spent outside of the cage, hours of sleep, lack of enrichments, length of ownership, lives with others parrots, position of the cage, bird sex, no toys/only one toy in the cage, inability to fly, other non-bird companion animals (protective), out of the cage for more than 8h, owner type (Shelter, Woman, Man, Family), being acquired from a pet shop, rearing method, being rescued or rehomed, separation anxiety, species, sprayed with water daily, time of human/bird interaction per day | *Agapornis*, *Amazona*, *Cacatua*, *Guaruba*, "Multiple", *Psittacus* | (2, 7, 8, 13, 15, 23, 49-60) |
| **Social behaviours** | Aggressive | 18 (3.14%) | Total aggressive behaviours (F, D), female/male aggressive behaviours (F), aggressive behaviours toward non-mates (F), wins/losses after agonistic interactions (F), number of chicks killed | Dominance rank, agonistic interactions, partner/non-partner affiliation, inappropriate handling, bird sex, mate/non-mate, lack of social interactions | *Amazona*, *Melopsittacus*, *Myiopsitta*, *Nymphicus* | (20, 61-66) |
| | Allopreen | 12 (2.09%) | Male/female allopreen (F), male/female allopreen solicitation (F), total allopreening (D, PP, P) | Sex of the receiver, cage size, feather damaging behaviours, lack of enrichments, type of physical enrichment, preference for partner | *Amazona*, *Ara*, *Melopsittacus*, *Nymphicus*, *Pyrrhura* | (11, 19, 21, 34, 61, 62) |
| | Sexual behaviours | 19 (3.31%) | Female/male courting (D, F), intra-pair interaction (D), number of approaches toward mates, synchronicity, female/male copulation (F), female solicits copulation (F), sexually active (U), sexual behaviours (P) | Positive correlated with feather damaging behaviours, cage size, lack of social interaction, bird sex, comparison between mate and non-mate, lack of physical, foraging, and cognitive enrichments, rearing methods, removed from the nest before 2 week/hatched from egg incubator | *Ara*, *Melopsittacus*, *Nymphicus*, *Psittacus*, *Pyrrhura* | (2, 4, 14, 34, 62) |
| | Social dynamics | 21 (3.66%) | Reaction towards other individuals (F), coo-feeding (F), food steeled (F, number of parrots), social interactions (PP), time spent next to each other, approach unfamiliar bird (L), dominance rank, physical distance between subjects (score) | Inappropriate handling, kinship, age, aggressiveness, time of day, bird sex, lack of enrichment, social isolation, cage size, lack of mate | *Ara*, *Melopsittacus*, *Nymphicus* | (19, 20, 30, 61, 62, 65, 67) |
| | Vocalizations | 23 (4.01%) | Calm vocalization (F), singing (D, F), vocalization (D, F, P), playback response (F) | Affiliation with other birds, negative correlation with feather damaging behaviours, positive correlation with accepting food from humans, lack of enrichments, pair housing, rearing method, having reduced flight ability, social deprivation, social isolation | *Amazona*, *Ara*, *Melopsittacus*, "Multiple", *Myiopsitta*, *Nymphicus*, *Pyrrhura* | (4, 9, 10, 21, 30, 34, 38, 64) |
| | Human-animal interaction | 34 (6.28%) | Attention bias, inappropriate handling, yawning after handling (L, F), response to unfamiliar/familiar handler (score), human aversion score, seeking behaviour toward humans (F), human-direct aggressiveness (yes/no), approach to humans (yes/no), food acceptance (yes/no, score), response to human contacts (score), resistance to being picked up (yes/no), latency to approach humans, begging once adult (yes/no), selective toward humans, tendency to anthropomorphize PCA axis*, vocalize during restrain (D, L, F), learned vocalization (F), long distance contact call (F) | Unfamiliar human presence, lack of human-animal interaction, inappropriate interaction, neophilia score, lack of neonatal handling, lack of enrichment, rearing method, mouth to beak feeding, human food consumption, owner gender, correlated with feather-damaging behaviours, lack of social interactions | *Amazona*, *Ara*, *Melopsittacus*, "Multiple", *Psittacus* | (2, 5, 9, 15, 16, 32, 37, 68-73) |
| | Facial and body displays | 12 (2.09%) | Crown, nape, lower/upper mandible, cheek feathers ruffling (scan), nape/crown feather height, erected crest (D), blushing around eyes | Correlation with positive human-parrot interaction, arousal level, social deprivation | *Ara*, *Cacatua*, *Nymphicus* | (67, 71, 73, 74) |

* Results obtained running a Principal Component Analysis

**Table S11**. Significant and not feasible outcome measures grouped by welfare dimensions and welfare categories. The table reports the type of unfeasible outcomes collected for all welfare categories, feasible risk factors associated with them, and all genera investigated with their corresponding studies. The outcomes' names are named as the authors reported them.

| Welfare dimension | Welfare categories | Types of unfeasible outcomes | Feasible risk factors | Genera | Studies |
|---|---|---|---|---|---|
| **Diseases and pathologic conditions** | Health | Presence of atherosclerosis, severity of atherosclerosis lesions, hepatic hemosiderosis, lipid accumulation lesions, prevalence of lipoid pneumonia, presence of ingluvioliths, health conditions, prevalence of hepatic lipidosis, presence of viral diseases | Age, rearing method, diet (processed vs seed based), species, fibres ingestion (bird sex, age), chicks artificial feeding method | *Amazona, "Multiple", Myiopsitta, Nymphicus, Psittacus* | (2, 46, 75-81) |
| **Body measurements** | Feathers Colour | Cheek, front and crown feathers chroma, front feathers hue, crown feathers luminance, cheek feathers structural colours | Inappropriate handling, unbalanced diet | *Platycercus* | (82) |
| **Physiological parameters** | Lipids-related | Cholesterol concentration, triglyceride concentration, high-density lipoprotein-cholesterol (HDL-C) concentration, low-density lipoprotein-cholesterol (LDL-C) concentration, ratio of total cholesterol and HDL-C, | Unbalanced diet, type of diet, bird sex, lack of exercise | *Amazona, Ara* | (46-48) |
| | Immune System-related | Humoral response to vaccination, delayed-type hypersensitivity (DTH) response, ratio of heterophils and lymphocytes, leukocytes, lymphocytes, and monocytes count | Lack of neonatal handling, unbalanced diet | *Amazona*, *Platycercus* | (46, 69, 82) |
| | Metabolic | Digestibility of crude fibres, crude proteins and dry matters, daily energy expenditure, malondialdehyde (MDA) concentration | Age, unbalanced diet, lack of exercise, type of diet, courtship feeding, weight, wing load (indirect measure of fat mass) | *Amazona*, Melopsittacus | (35, 36, 83, 84) |
| | Stress-related | Corticosterone excreta metabolites concentration, plasma corticosterone, cortisol excreta metabolites concentration, | Age, agonistic interactions, artificial light at night, dominance rank, bird sex, positively correlation with feather damaging behaviour, positive correlation with foraging time, inappropriate handling, lack of neonatal handling, living conditions (wild, zoo, breeding centre, companions), social isolation, negative correlation with locomotor behaviours before implementing enrichment, negative correlation with object interaction before implementing enrichment, being wild caught | *Agapornis, Amazona*, *Ara*, *Melopsittacus*, "Multiple", *Nymphicus*, *Platycercus, Psittacus* | (10, 20, 41, 60, 66, 69, 82, 85-88) |
| | Vitamin D-related | Calcifediol concentration, plasma vitamin-D concentration, plasma Ca+ concentration, plasma Mg+ concentration, | Indoor housing, lack of UV light | *Amazona* | (89, 90) |
| | Others | DNA damage, telomere length, basal glucose concentration, respiration rate, aortic pressure gradient and speed, hemoglobulin concentration, aspartate amino transferase concentration, glucose concentration, | Age, unbalanced diet, type of diet, lack of neonatal handling, social isolation, weight, activity level | *Amazona*, *Ara*, *Melopsittacus*, *Psittacus* | (46, 47, 68, 69, 84, 91, 92) |

**Table 12**. Significant and feasible outcomes measures reported in companion parrots specifically. The table shows the types of outcomes collected, the number of outcomes, their corresponding risk factors, the genera and the intervention used in the studies. The names for outcome measures and risk factors are kept as the authors reported them.

| Welfare category | Types of outcomes | Number of outcomes (%) | Risk factors | Genera (number of outcomes) | Intervention (number of outcomes) |
|---|---|---|---|---|---|
| **Indirect measures of feather damaging behaviours** | Presence/absence of feather damages, plumage score | 51 (75%) | Due to high number, see Table S13 | *Agapornis* (3), *Cacatua* (9), *Psittacus* (10), Multiple (29) | Video-Analysis (5), Clomipramine treatment (2), Questionnaire (39), Check-up at veterinary clinic (3) |
| **Fear-related** | Phobic behaviour | 1 (1.47%) | Being wild caught | *Psittacus* | Questionnaire |
| | | 2 (2.94%) | Perches lower than eye level | | |
| | | | Small cage (max. 80 cm x 100 cm x 120 cm) | | |
| **Foraging** | Contra-freeloading as foraging enrichment | 1 (1.47%) | Individuality | *Psittacus* | Contra-freeloading test |
| **Human-Animal Interaction** | Anthropomorphising | 1 (1.47%) | Female ownership | Multiple | Questionnaire |
| | Begging once adult | 3 (4.6%) | Hand-rearing | *Psittacus* | |
| | | | Human mouth to beak feeding | | |
| | | | Human leftover consumption | | |
| | Selective towards humans | 1 (1.47%) | Hand-rearing | | |
| **Stereotypies** | Multiple Stereotypies | 1 (1.47%) | Inappropriate diet | *Psittacus* | Questionnaire |
| | | 1 (1.47%) | Only manufactured perches in the cage | | |
| | | 1 (1.47%) | Removed from the nest before 5 weeks | | |
| | Whole Body Stereotypies | 1 (1.47%) | Positive correlated with brain volume (species) | Multiple | Questionnaire |
| | Oral Stereotypies | 1 (1.47%) | | | |
| **Sexual behaviours** | Sexual activity | 1 (1.54%) | Removed from the nest before 2 weeks or hatched from egg incubator | *Psittacus* | Questionnaire |
| **Self-care** | Preening | 1 (1.54%) | Hand-rearing | *Psittacus* | Questionnaire |
| | | 1 (1.54%) | Sold before the end of weaning | | |

*Hardly any fruit at all/only few different sorts/no separate bowl for fruit or leftovers/food for human consumption or little fruit, inappropriate seed mixture and no protein supply.

#### Outcomes related to feather damaging behaviours

From the 690 significant results, 70 outcomes, collected from 19 different studies, were related to feather damaging behaviours. Most of the outcomes were collected from companion animals (n=51), followed by parrots kept in laboratories (n=11), shelter parrots (n=6) and parrots kept in breeding and rehab centres (both n=1). Most of the outcomes (n=22) were associated with risk factors belonging to demographic characteristics such as age, sex, and species (Table S13). The lack of enrichment opportunities (14 outcomes) and factors related to the human-animal relationship (15 outcomes) turned out to also be common potential risk factors for feather-damaging behaviour (Table S13).

**Table S13**. Outcomes and studies related to feather damaging behaviours and corresponding risk factors.

| Category | Risk factor | Outcomes | Number of Studies | Genera | Living condition |
|---|---|---|---|---|---|
| **Ease of Movement** | Inability to fly | 1 | 1 | *Psittacus* | Companion |
| **Demographic** | Increasing age | 6 | 4 | *Agapornis*, Multiple, *Psittacus* | Companion |
| | Heritability | 2 | 1 | *Amazona* | Lab |
| | Sex | 3 | 3 | *Amazona*, *Cacatua*, Multiple | Companion (2), Lab (1) |
| | Being rescued | 1 | 1 | Multiple | Companion |
| | Species | 9 | 3 | Multiple | Companion |
| | Diet needing extensive handling (species) | 1 | 1 | Multiple | Companion |
| **Enrichment** | Lack of chewable devices | 1 | 1 | Multiple | Companion |
| | Lack of foraging devices | 1 | 1 | Multiple | Companion |
| | Hour spent outside of the cage | 1 | 1 | Multiple | Companion |
| | Lack of foraging + physical Enrichment | 2 | 1 | *Amazona* | Lab |
| | Lack of foraging + human and physical enrichment | 4 | 1 | *Amazona* | Lab |
| | Lack foraging Enrichment | 3 | 1 | *Psittacus* | Shelter |
| | No toys/only one toy in the cage | 1 | 1 | *Psittacus* | Companion |
| | Out of the cage > 8h | 1 | 1 | Multiple | Companion |
| **Good Human and Animal relationship** | Number of vacations per year | 2 | 1 | *Cacatua, Psittacus* | Companion |
| | Command training | 1 | 1 | *Cacatua* | Companion |
| | Length of ownership | 1 | 1 | *Psittacus* | Companion |
| | Owner type (shelter, woman, man, family) | 1 | 1 | Multiple | Companion |
| | Bought from a pet shop | 1 | 1 | *Cacatua* | Companion |
| | Rearing method | 6 | 4 | *Amazona*, *Agapornis* Multiple, *Psittacus* | Companion (4), Lab (1) |
| | Separation anxiety | 2 | 2 | *Agapornis*, Multiple | Companion |
| | Time of human-parrot interaction per day | 1 | 1 | Multiple | Companion |
| **Good Housing** | Cage volume > 2 m³ | 1 | 1 | *Cacatua* | Companion |
| | Distance of the cage from the door | 1 | 1 | *Amazona* | Lab |
| | Location of the cage against >= 1m wall | 1 | 1 | *Cacatua* | Companion |
| **Enrichment+Personality** | Lack of enrichments + neuroticism score | 1 | 1 | *Psittacus* | Lab |
| **Maintenance** | Hours of sleep | 1 | 1 | *Psittacus* | Companion |
| | Sprayed with water daily | 1 | 1 | *Cacatua* | Companion |
| **Good Feeding** | Fed with only seed or pellet | 1 | 1 | Multiple | Companion |
| **Personality** | Coping style (proactive) | 3 | 1 | *Psittacus* | Shelter |
| **Physiological** | Adrenocortical activity | 1 | 1 | *Guaruba* | Breeding Centre |
| **Social** | Lives with other parrots | 1 | 1 | Multiple | Companion |
| | Lives without other non-bird companion animals | 1 | 1 | Multiple | Companion |
